# PEX1^G843D^ remains functional in peroxisome biogenesis but is rapidly degraded by the proteasome

**DOI:** 10.1101/2024.12.10.627778

**Authors:** Connor J. Sheedy, Soham P. Chowdhury, Bashir A. Ali, Julia Miyamoto, Eric Z. Pang, Julien Bacal, Katherine U. Tavasoli, Chris D. Richardson, Brooke M. Gardner

## Abstract

The PEX1/PEX6 AAA-ATPase is required for the biogenesis and maintenance of peroxisomes. Mutations in *HsPEX1* and *HsPEX6* disrupt peroxisomal matrix protein import and are the leading cause of Peroxisome Biogenesis Disorders (PBDs). The most common disease-causing mutation in PEX1 is the *Hs*PEX1^G843D^ allele, which results in a reduction of peroxisomal protein import. Here we demonstrate that *in vitro* the homologous yeast mutant, *Sc*Pex1^G700D^, reduces the stability of Pex1’s active D2 ATPase domain and impairs assembly with Pex6, but can still form an active AAA-ATPase motor. *In vivo*, *Sc*Pex1^G700D^ exhibits only a slight defect in peroxisome import. We generated model human *Hs*PEX1^G843D^ cell lines and show that PEX1^G843D^ is rapidly degraded by the proteasome, but that induced overexpression of PEX1^G843D^ can restore peroxisome import. Additionally, we found that the G843D mutation reduces PEX1’s affinity for PEX6, and that impaired assembly is sufficient to induce degradation of PEX1^WT^. Lastly, we found that fusing a deubiquitinase to PEX1^G843D^ significantly hinders its degradation in mammalian cells. Altogether, our findings suggest a novel regulatory mechanism for PEX1/PEX6 hexamer assembly and highlight the potential of protein stabilization as a therapeutic strategy for PBDs arising from the G843D mutation and other PEX1 hypomorphs.

## Introduction

Peroxisomes are membrane-bound organelles that are present in most eukaryotic cells (1). They harbor a dense matrix of enzymes to carry out unique metabolic reactions such as the beta-oxidation of very long chain fatty acids, the detoxification of reactive oxygen species (ROS), and the biosynthesis of plasmalogens contained in myelin (2–5). Peroxisomes display a remarkable diversity of function: they participate in metabolic crosstalk with other organelles, innate immunity and antiviral signaling, and tissue-specific functions (6–9). This adaptability suggests that cells can rapidly tune their peroxisome function according to environmental cues or pathogens. Owing to their roles in lipid metabolism and ROS detoxification, peroxisomes are also essential for proper brain development and neurological function (10, 11).

The formation and maintenance of peroxisomes relies on ∼35 peroxins encoded by *PEX* genes (12). Mutations in at least 13 different *PEX* genes cause peroxisome biogenesis disorders (PBDs) that are characterized by widespread metabolic dysfunction (13). PBDs are divided into the broad group of Zellweger spectrum disorders (PBD-ZSD) and the distinct rhizomelic chondrodysplasia punctata type 1 (14). Patients presenting with PBD-ZSD display a wide breadth of phenotype severity, ranging from developmental abnormalities and early infant mortality, to progressive sensorineural hearing loss and retinal degeneration (15). Due to the complex phenotypes presented by peroxisomal dysfunction, and the many genes associated with regulation of the organelle, a number of other disorders such as Usher syndrome are often misdiagnosed and later revealed to be PBDs through deep sequencing approaches (16). PBD-ZSD patients typically exhibit progressive deafness and retinopathy, but certain preventative measures such as diet restriction and avoidance of hepatotoxic substances have been shown to ameliorate symptoms. Therefore, strategies to improve peroxisomal protein targeting may be of broad benefit to PBD-ZSD patients, as well as the numerous other conditions in which peroxisome function is dysregulated.

The majority of PBD-ZSDs are caused by mutations in either *PEX1* or its binding partner *PEX6* (58.9% and 15.9%, respectively) (13). PEX1 and PEX6 assemble into a heterohexameric mechanoenzyme in the family of type-II AAA-ATPases, which contain two ATPase rings (17, 18). Proteins in this family use the energy of ATP hydrolysis to unfold substrates by processive threading or to disassemble protein complexes through large conformational changes, as exemplified by ClpB and human NSF, respectively (19, 20). The ATPase activity of PEX1/PEX6 is thought to drive the import of matrix proteins into the peroxisome by extracting the peroxisome targeting signal receptor PEX5 from the peroxisome membrane (21–24). In the absence of PEX1/PEX6 function, PEX5 accumulates in the peroxisomal membrane (22), matrix protein import is impaired (25), and peroxisomes are targeted for peroxisome-specific autophagy (26, 27).

The most prevalent disease-causing mutation in PEX1, accounting for nearly 45% of observed cases, is a missense mutation converting glycine 843 to an aspartic acid (G843D) (28). Patients with the G843D mutation display milder phenotypes in the Zellweger spectrum and multiple lines of evidence suggest that PEX1^G843D^ is a hypomorphic allele (29). Patient fibroblasts homozygous for the G843D mutation exhibit a reduction of peroxisomal matrix protein import that can be partially restored by culturing cells at 30°C (29, 30), by transfection with a plasmid expressing PEX1^G843D^ (31), and by overexpression of the PEX1 binding partner PEX6 (32). The G843D mutation reduces the level of PEX1 proteins in cells, a change that is not explained by a concomitant decrease in mRNA transcript levels (29, 33). Treatments that ameliorate the import defect in PEX1^G843D^ fibroblasts, such as growth at a reduced temperature of 30°C, also increase PEX1 protein levels (29). Thus, the prevailing model in the field is that the G843D mutation causes a folding defect in PEX1 that destabilizes PEX1 and disrupts its interaction with PEX6 (29). However, it has not yet been shown how PEX1^G843D^ is degraded or how the G843D mutation impacts motor function. Mutations in PEX1 and PEX6 have been shown to induce peroxisome specific autophagy (27), and there are conflicting reports regarding whether the inhibition of autophagy can improve peroxisome import in PEX1 hypomorphs (26, 34).

Recent advances in protein prediction and studies with the functionally homologous yeast Pex1/Pex6 complex suggest high structural conservation between experimentally determined yeast structures (35, 36) and AlphaFold-predicted human structures (37). These comparisons have allowed more accurate mapping of disease mutations and their proposed effects on PEX1/PEX6 function (38, 39). Similar to yeast Pex1/Pex6, human PEX1 and PEX6 each contain two N-terminal domains, N1 and N2, thought to be involved in substrate binding, and two ATPase domains, D1 and D2. The inactive D1 ATPase ring supports hexamerization, while the active D2 ATPase ring hydrolyzes ATP and drives substrate processing through conformational changes driven by ATP-hydrolysis. The G843D mutation occurs at the transition between the D1-D2 linker and the beginning of the D2 ATPase domain. Given the location of the mutation and the well conserved nature of the glycine in this position, it is feasible that G843D alters D2 ATPase domain folding, ATPase activity, and/or assembly with PEX6. In this study, we set out to dissect how the G843D mutant impairs PEX1/PEX6 activity.

We utilize *in vitro* biochemistry techniques to demonstrate that the homologous G843D variant in yeast PEX1, *Sc*Pex1^G700D^, reduces the stability of the isolated D2 ATPase domain. We further find that the purified *Sc*Pex1^G700D^/*Sc*Pex6 hexamer displays residual ATPase and unfoldase activity, providing evidence for cooperative complex assembly in the presence of ATP. We then generate disease-model human cell lines and demonstrate that PEX1^G843D^ is rapidly degraded by the ubiquitin proteasome network, but otherwise capable of supporting peroxisome function *in vivo*. We additionally demonstrate that PEX1^G843D^ protein levels can be modestly stabilized by the knockdown of specific E3 ligases, and dramatically stabilized in mammalian cells when fused to a deubiquitinase, OTUB1, suggesting that protein stabilization may be a promising therapeutic avenue. Finally, we find evidence that the stability of wildtype PEX1 is reduced in the absence of PEX6, indicating that orphan protein quality control mechanisms may fine tune the stoichiometry of the peroxisomal import machinery.

## Results

### Yeast Pex1^G700D^/Pex6 is a functional ATP unfoldase

The pathogenic PEX1^G843D^ mutation replaces a glycine in the linker preceding the D2 ATPase domain with aspartic acid (Figure 1A-C, Figure S1). Previous work has shown that recovery of peroxisome import in PEX1^G843D^ cells occurs at lower temperatures and upon addition of small molecule chaperones, both suggesting a defect in the folding of PEX1 caused by the G843D mutation (29, 30, 40, 41). As the glycine at this position is conserved from humans to yeast (Figure 1D), and the predicted structure of this region of *Hs*PEX1 is highly similar to that of *Sc*Pex1 (Figure 1B-1C, Figure S1), we set out to assess how this mutation alters the folding of the yeast Pex1 D2 ATPase domain. We purified the isolated D2 ATPase domain as an MBP fusion protein and measured domain stability using a SYPRO-Orange thermal melt assay. We observed two unfolding events for the MBP-*Sc*Pex1^WT^ D2 ATPase fusion. One unfolding event observed at T_m_ = 52°C was not sensitive to ATP and is presumably the MBP domain. Another unfolding event, presumably corresponding to the D2 ATPase domain, was sensitive to the concentration of ATP: In the absence of ATP, unfolding was observed at T_m_ = 23.67°C, while in the presence of 10 mM ATP, unfolding was observed at Tm = 38.8°C (Figure 1E). In comparison, we found that the G700D mutation reduced the melting temperature of the D2 ATPase domain to 27.5°C from 38.8°C for *Sc*Pex1^WT^ in the presence of 10 mM ATP (Figure 1E). We were unable to observe temperature induced unfolding of *Sc*Pex1^G700D^ at lower concentrations of ATP. We therefore conclude that G700D destabilizes the isolated *Sc*Pex1 D2 ATPase domain, but that folding can be partially stabilized by nucleotide binding.

**Figure 1:**
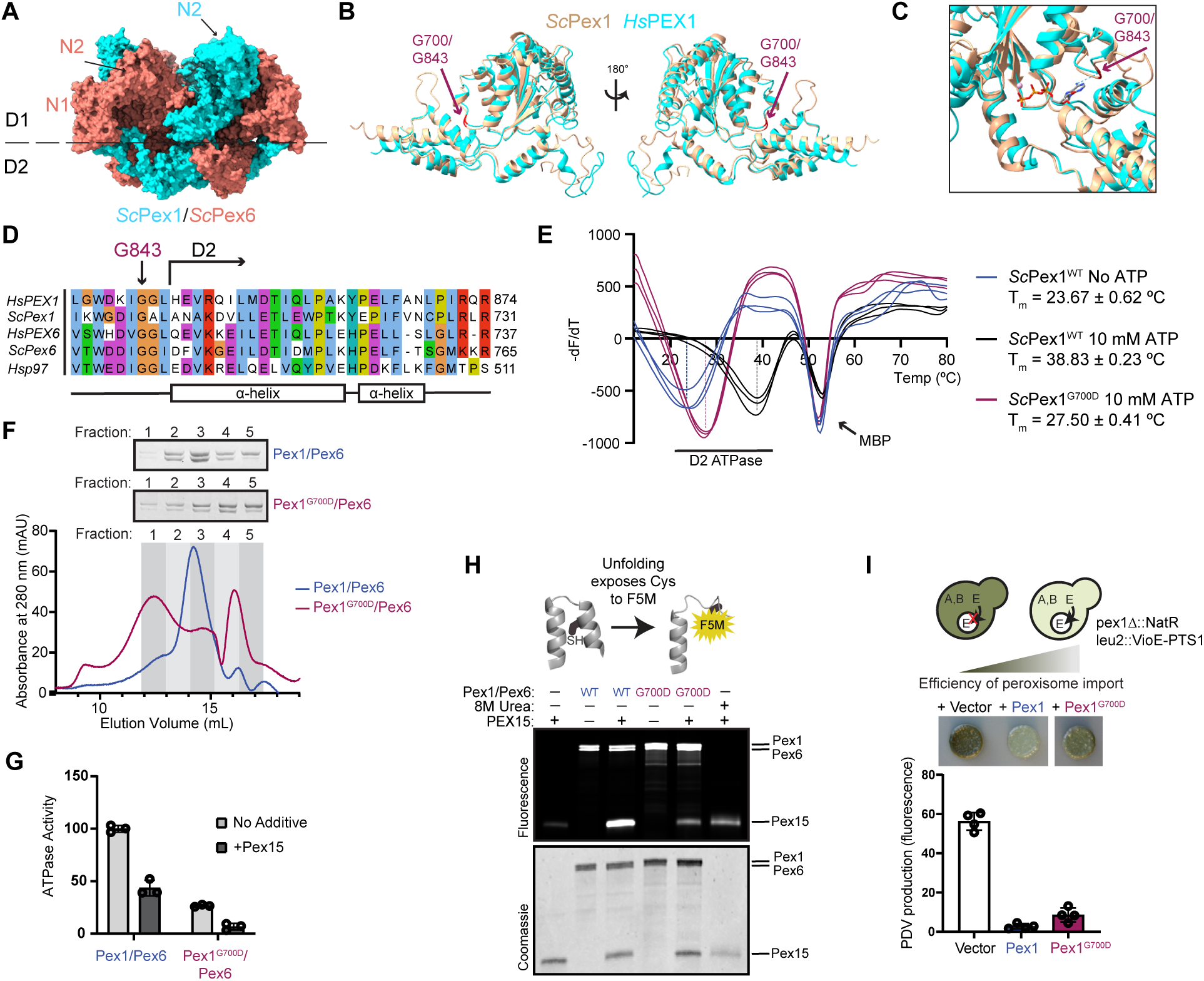
Yeast Pex1^G700D^ is a partially active unfoldase in vitro but supports peroxisome import in vivo. A) Structure and domain architecture of heterohexameric *S. cerevisiae* Pex1/Pex6. PDB:8U0V. B) Structural alignment of AlphaFold2 predictions human (cyan) and yeast (tan) Pex1 D2 ATPase domains highlighting G843 and G700 residues (red)(37). C) As in B, focus on ATP binding pocket. ATP is modeled from AlphaFold2 structures based on alignment with human p97 (PDB: 7LN5) (64). D) Sequence alignment of the G843 position in *Hs*PEX1 with *Sc*Pex1, *Hs*PEX6, *Sc*Pex6, and *Hs*p97. E) SYPRO-Orange thermal melt assays of the MBP-D2 ATPase fusion. The plot shows the inverse derivative of raw RFU values for each replicate. Means are shown with SD of N=3 replicates. F) Representative chromatograms of recombinantly purified *Sc*Pex1^G700D^/*Sc*Pex6 and *Sc*Pex1^WT^/*Sc*Pex6 on a Superose 6 Increase and Coomassie-stained SDS-PAGE of the fractions. G) ATPase assay using *S. cerevisiae* Pex1/Pex6 and Pex1^G700D^/Pex6 complexes and *Sc*Pex15 (1-309) recombinantly purified from *E. coli*. Means with SD are shown from N=3 replicates. H) Fluoroscein-5-maleimide (F5M) labeling experiments indicating *Sc*Pex1/*Sc*Pex6-mediated unfolding of *Sc*Pex15 (1-309). *Sc*Pex15 contains buried cysteines that are labeled by F5M upon unfolding. Urea is a control for unfolding of *Sc*Pex15 without motor present. I) Colorimetric assay for peroxisome import to measure complementation in a *Δpex1* deletion strain by *Sc*Pex1^G700D^ and *Sc*Pex1^WT^ with plate phenotype and quantification of PDV pigment production. Means with SD are shown from N=4 replicates.

To test the functionality of the G700D mutant in the context of the heterohexameric Pex1/Pex6 complex, we purified recombinant *Sc*Pex1^G700D^/*Sc*Pex6 complexes. Using size-exclusion chromatography, we found that purified *Sc*Pex1^G700D^/*Sc*Pex6 reduced the proportion of Pex1/Pex6 in the hexameric fractions compared to the wild-type complex (Figure 1F, fractions 2 and 3), and increased the proportion of slower migrating species, presumably Pex1 and Pex6 monomer or dimers (Figure 1F, fractions 4 and 5, Figure S2A). This shift indicates an increase in the rate of hexamer disassembly or an assembly defect conferred by the G700D mutation. Despite this assembly defect, the *Sc*Pex1^G700D^/*Sc*Pex6 retained ∼30% ATPase activity compared to wild-type *Sc*Pex1/*Sc*Pex6 (Figure 1G), and was also inhibited by the cytosolic domain of *Sc*Pex15 (Figure 1G, Figure S2B) (24). Wild-type *Sc*Pex1/*Sc*Pex6 was previously shown to unfold truncated *Sc*Pex15 *in vitro* in a manner that depended on both active Pex1 and Pex6 D2 ATPase domains (24). Using a maleimide labeling assay for unfolding, in which we measure the labeling efficiency of buried cysteine residues as they become exposed, we found that the *Sc*Pex1^G700D^/*Sc*Pex6 complex unfolds truncated *Sc*Pex15, albeit with reduced efficiency compared to wild-type *Sc*Pex1/*Sc*Pex6 (Figure 1H). These results suggest that although *Sc*Pex1^G700D^ assembly with *Sc*Pex6 is impaired, it retains the ability to form a functional AAA-ATPase that is capable of unfolding substrates.

To assess the *in vivo* effects of the mutation on Pex1/Pex6 function in *S. cerevisiae*, we performed a complementation experiment in a *Δpex1* deletion strain harboring a colorimetric peroxisome import reporter. This strain is engineered to compartmentalize a key enzyme in the prodeoxyviolacein pathway to the peroxisome, such that when peroxisome import is impaired, the yeast produce a green pigment (Figure 1I) (42). Re-introduction of *Sc*Pex1^G700D^ in a *Δpex1* deletion strain reduced green pigment formation nearly to the same levels as *Sc*Pex1^WT^, indicating that the *Sc*Pex1^G700D^ complements the *Δpex1* deletion with a mild defect (Figure 1I). The observed import defect with the G700D mutation is therefore consistent with an import defect in patient fibroblasts expressing the G843D mutation (29). Taken together, we conclude that the G700D mutation partially compromises the folding of the Pex1 D2 ATPase domain and the assembly of hexamer with Pex6, but in the context of full length Pex1, Pex6 binding partner, and high ATP concentrations, the *Sc*Pex1^G700D^ D2 ATPase domain can support peroxisome import function *in vivo*.

### The G843D mutation disrupts peroxisome import and decreases PEX1 protein levels in patient fibroblasts and in model cell lines

Previous work in patient fibroblasts carrying the G843D mutation showed a strong reduction in the levels of PEX1 protein that was not explained by a concomitant reduction in mRNA levels. We confirmed markedly decreased PEX1 protein levels in immortalized and primary patient fibroblasts with the G843D mutation (Figure 2A, Figure S3A). The G843D fibroblasts had a modest, but statistically insignificant reduction in *PEX1* mRNA transcript levels and a slight increase in *PEX6* mRNA transcripts (Figure 2B). As expected, PEX1^G843D/G843D^ cells presented a peroxisome import defect seen by significantly reduced co-localization of peroxisomal membrane protein PMP70 and peroxisomal matrix enzyme catalase (Figure 2C). Due to the confounding effects of donor-to-donor variability in patient derived cell lines, as well as the limitations of genetic tractability, we sought to create PEX1^G843D^ model cell lines to better isolate our study of the effects of PEX1^G843D^ (28, 43).

**Figure 2.**
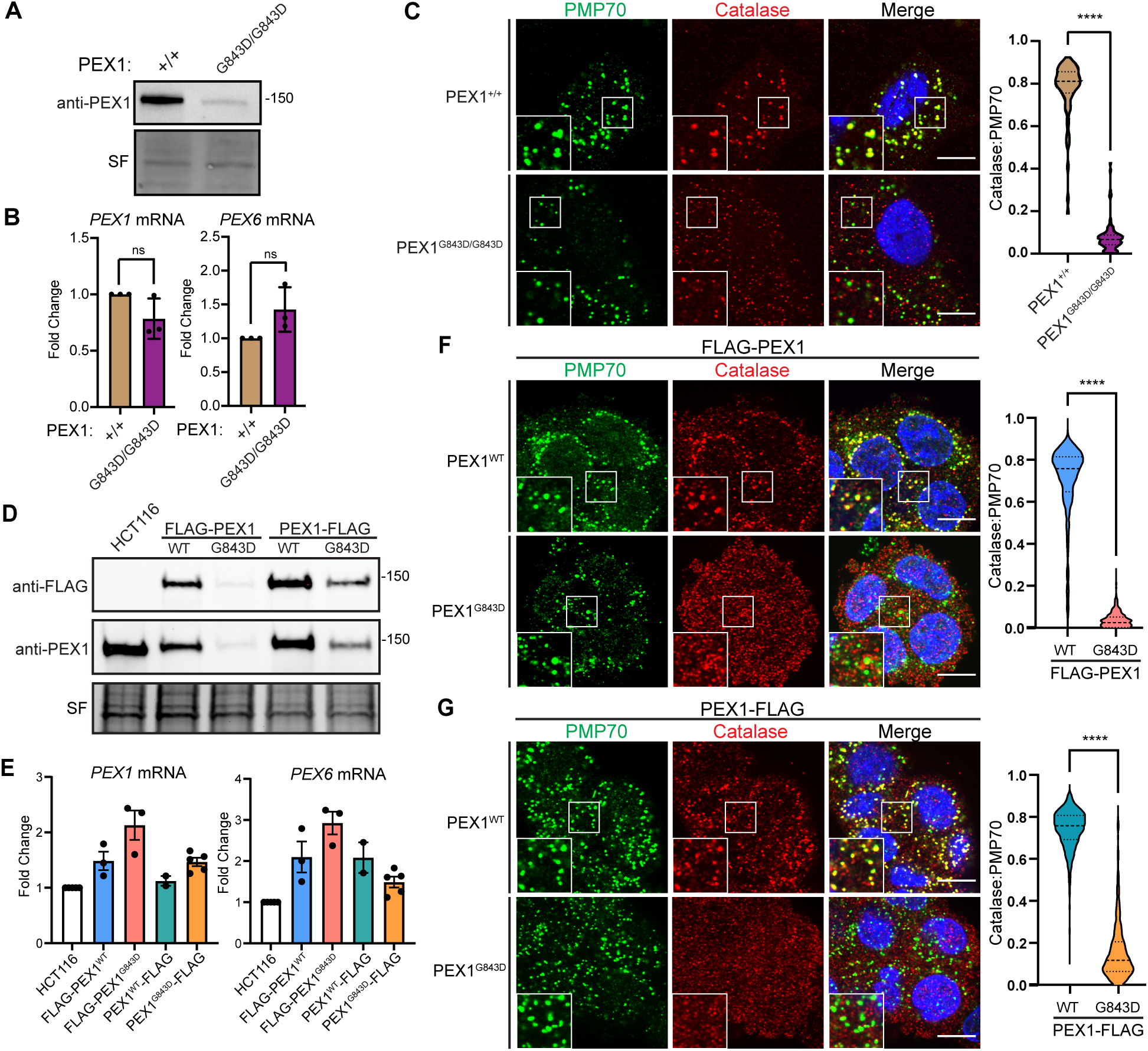
Disease model HCT116 PEX1^G843D^ cell lines recapitulate PBD patient fibroblast phenotypes. A) Representative immunoblots of immortalized patient fibroblasts show decreased levels of PEX1^G843D^ (PEX1 = 142 kDa, SF = StainFree total protein). B) RT-qPCR in immortalized patient fibroblasts with genotypes of PEX1^+/+^ or PEX1^G843D/G843D^ indicate no significant change of *PEX1* and *PEX6* mRNAs. Means are shown with SD from N = 3 replicates, unpaired t-test with Welch’s correction. C) Representative images and quantification of PMP70/catalase co-localization from immunofluorescence microscopy of immortalized patient fibroblasts. Panels show PMP70 (green), catalase (red), and merged images with DAPI (blue) and insets show the colocalization of puncta. Scale bar = 10 μm. Unpaired t-test with Welch’s correction, p-value < 0.0001. D) Representative immunoblots for anti-FLAG or anti-PEX1 in the HCT116 CRISPR-Cas9 edited clonal FLAG-PEX1/FLAG-PEX1, FLAG-PEX1^G843D^/FLAG-PEX1^G843D^, PEX1-FLAG/PEX1, and PEX1^G843D^-FLAG/PEX1^G843D^ cell lines. E) RT-qPCR in HCT116 FLAG-tagged PEX1 clonal cells indicate a slight increase in both *PEX1* and *PEX6* mRNA levels compared to unedited HCT116 cells. N ≥ 2. F) As in (C) with HCT116 FLAG-PEX1/FLAG-PEX1 and FLAG-PEX1^G843D^/FLAG-PEX1^G843D^. Scale bar = 10 μm. Unpaired t-test with Welch’s correction, p-value < 0.0001. G) As in (C) with HCT116 PEX1-FLAG/PEX1, and PEX1^G843D^-FLAG/PEX1^G843D^ cell lines. Scale bar = 10 μm. Unpaired t-test with Welch’s correction, p-value < 0.0001.

To create PEX1^G843D^ model cell lines, we first used Cas9 gene editing to create clonal homozygous N-terminal FLAG-PEX1 and heterozygous C-terminal PEX1-FLAG cell lines in human male colorectal carcinoma HCT116 cells. Into these two FLAG-tagged cell lines we used Cas9 gene editing to simultaneously introduce the G843D mutation and a silent mutation for a ClaI restriction enzyme cleavage site (Figure S3B). We then screened clonal cells by ClaI digestion of PCR amplicons of the PEX1 gene from genomic DNA to identify clonal populations homozygous for PEX1^G843D^ (Figure S3C).

In both FLAG-tagged cell lines, the introduction of the G843D allele significantly reduced PEX1 protein levels (Figure 2D), recapitulating the reduced PEX1 abundance observed in patient fibroblasts. Analysis of *PEX1* and *PEX6* mRNA levels by RT-qPCR showed that introduction of the G843D mutation did not significantly alter *PEX1* or *PEX6* mRNA levels, similar to patient fibroblasts (Figure 2E). Both FLAG-PEX1^G843D^ and PEX1^G843D^-FLAG cell lines exhibited reduced co-localization of peroxisomal membrane protein PMP70 and peroxisomal matrix enzyme catalase, recapitulating the import defect observed in patient fibroblasts (Figure 2F-2G). Overall, the HCT116 cell lines with epitope tagged PEX1^G843D^ phenotypically resemble PEX1^G843D^ patient fibroblasts and provide a useful tool for interrogating the molecular effects of the G843D mutation.

### PEX1^G843D^ is rapidly degraded by the proteasome

The observed reduction of PEX1 protein without reduction of *PEX1* mRNA suggests that the G843D mutation either reduces PEX1 mRNA translation or PEX1 protein stability. To determine if the reduction of PEX1^G843D^ protein levels arises from increased degradation, we performed *in vivo* cycloheximide chase experiments to compare the rate of degradation of FLAG-PEX1 and FLAG-PEX1^G843D^ homozygous HCT116 cell lines when uncoupled from the rate of translation. FLAG-PEX1 was stable over 24 hours after translation was stalled with cycloheximide, whereas FLAG-PEX1^G843D^ was rapidly degraded with a half-life of 3.12 ± 1.17 hours (Figure 3A-3B). Others have observed increased autophagic degradation of peroxisomes, or pexophagy, in patient cells with the G843D mutation, raising the question if pexophagy is responsible for PEX1^G843D^ degradation (26, 28, 29). We performed a similar experiment to test this hypothesis by culturing cells with either cycloheximide, or cycloheximide along with bafilomycin A1, an inhibitor of autophagosome and lysosome fusion, and therefore an inhibitor of autophagic degradation. The addition of autophagy inhibitor bafilomycin did not stabilize FLAG-PEX1^G843D^ in our cycloheximide chase assay indicating that elevated autophagy activation was not the degradative mechanism for FLAG-PEX1^G843D^ (Figure 3C-3D). We note that the autophagy inhibitor bafilomycin A1 resulted in an accumulation of LC3-II as expected, indicating that the treatment was effective at inhibiting autophagy. We then interrogated whether degradation was dependent on the proteasome. We found that the addition of the irreversible proteasome inhibitor carfilzomib greatly attenuated PEX1^G843D^ degradation with the half-life increasing from 3.12 ± 1.17 hours to greater than the 8-hour duration of the experiment (Figure 3E-3F). Thus, the proteasome degrades PEX1 harboring the G843D mutation.

**Figure 3:**
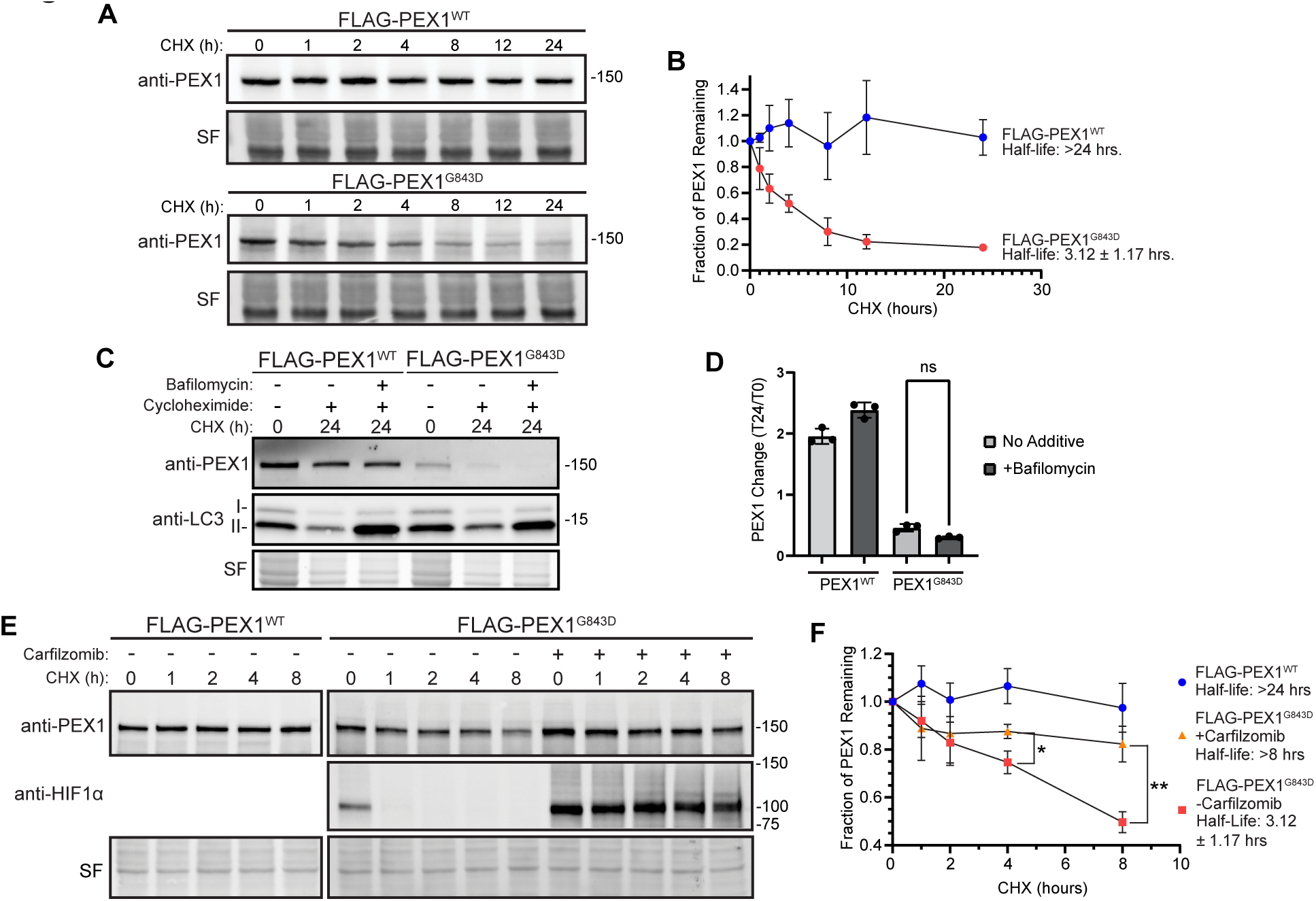
PEX1^G843D^ is rapidly degraded by the proteasome and not autophagy. A) Representative immunoblots of cycloheximide (CHX) chase assay in FLAG-PEX1^WT^ and FLAG-PEX1^G843D^ clonal cell lines (PEX1 = 142 kDa, SF = StainFree total protein). B) Quantification of CHX-chase assays shown in (A). Means are shown with SD from N=4 replicates for each timepoint. C) Representative immunoblots of PEX1 and LC3 in CHX-chase assay with the autophagy inhibitor Bafilomycin A1 (PEX1 = 142 kDa, LC3-I = 16-18 kDa, LC3-II = 14-16 kDa, SF = StainFree total protein). D) Quantification of autophagy inhibition CHX-chase assays shown in (C). Means are shown from N=3 replicates for each timepoint. Unpaired t-test with Welch’s correction. E) Representative immunoblots of PEX1 and HIF1α in CHX-chase assay with proteasome inhibitor Carfilzomib (PEX1 = 142 kDa, HIF1α = ∼100 kDa, SF = StainFree total protein). F) Quantification of proteasome inhibition CHX-chase assays shown in (E). Means are shown from N=3 replicates for each timepoint. Unpaired t-test with Welch’s correction. P-values: * = 0.02259, ** = 0.005724.

### PEX1^G843D^ can support peroxisome import function

Our data in yeast showed that the homologous variant *Sc*Pex1^G700D^ could support peroxisome import *in vivo*, suggesting that PEX1^G843D^ may be functional if it were not targeted for degradation. To assess whether PEX1^G843D^ in human cells is similarly capable of supporting peroxisome import function, we used lentiviral transduction to stably introduce TetOn-PEX1^WT^ or TetOn-PEX1^G843D^ constructs into our FLAG-PEX1^G843D^ cell line to allow for tunable overexpression with doxycycline (Figure 4A). The doxycycline-induced overexpression of PEX1^WT^ or PEX1^G843D^ was sufficient to restore PEX1 protein levels (Figure 4B). We note that cells required higher concentrations of doxycycline to restore PEX1^G843D^ to wild-type levels. To assess whether peroxisome import function was restored by doxycycline induced overexpression, we used the proteolytic processing of SCP2 in the peroxisome as a readout. Full length SCP2 (58 kDa) is cleaved to a smaller (46 kDa) active form upon peroxisomal import (44). Overexpression of PEX1^WT^ fully restored SCP2 processing whereas overexpression of PEX1^G843D^ strongly increased processing but was still partial compared to PEX1^WT^ (Figure 4B-4C). Furthermore, when peroxisomal import was assessed by immunofluorescence microscopy, we found that PEX1^G843D^ overexpression strongly rescued peroxisomal import as seen by colocalization of a peroxisomal matrix protein catalase and a membrane protein PMP70. Interestingly, the overexpression of PEX1^G843D^ produced a mosaic of rescue phenotypes compared to the full rescue with PEX1^WT^ overexpression (Figure 4D-4E). This observation highlights the strong propensity for PEX1^G843D^ to be degraded by the proteasome and suggests the existence of a “critical threshold” of PEX1^G843D^ that can fully rescue the import defect. Taken together, these data suggest that PBDs arising from the G843D mutation can be ameliorated by stabilizing PEX1^G843D^ protein levels.

**Figure 4:**
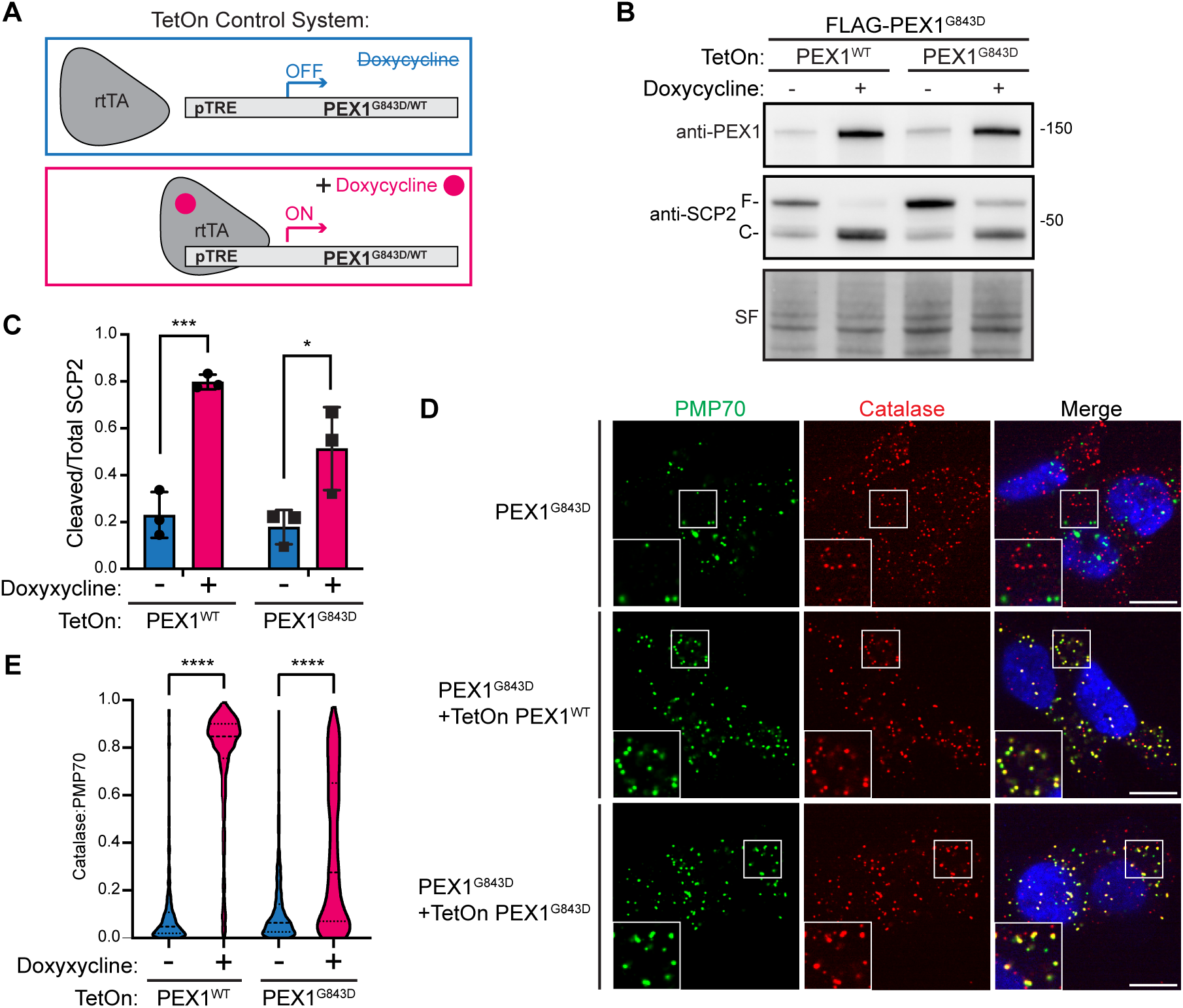
Overexpression of PEX1^G843D^ partially rescues peroxisome function. A) Schematic for the TetOn tunable gene expression constructs. Plasmid constructs contain a PEX1 variant expressed under Tet-responsive promoter (pTRE) and BFP-T2A-rtTA constitutively expressed under the hPGK promoter. Upon the addition of doxycycline, rtTA binds to pTRE to induce expression of PEX1 variants. B) Representative immunoblots of PEX1 and SCP2 with doxycycline induced overexpression of both PEX1^WT^ and PEX1^G843D^ in FLAG-PEX1^G843D^ HCT116s (PEX1 = 142 kDa, SF = StainFree total protein). Full-length SCP2 (SCP2-F, 58 kDa) is cleaved upon import into the peroxisome (SCP2-C, 46 kDa). C) Quantification of cleaved/total SCP2 upon induced expression of PEX1^WT^ or PEX1^G843D^ from replicates shown in (B). Means with SD are shown from N=3 replicates. Unpaired t-test, p-value summary: *** = 0.0007, * = 0.0379. D) Representative immunofluorescence microscopy images of catalase (red), PMP70 (green), and DAPI (blue) upon PEX1 overexpression. Insets show the colocalization of puncta. Scale bar = 10 µm. E) Quantification of catalase:PMP70 colocalization from experiments shown in (D). Unpaired t-test with Welch’s correction, all p-values < 0.0001.

### Assembly defect of the PEX1/PEX6 complex promotes degradation of PEX1

It has previously been shown that the G843D mutation reduces PEX1 binding to PEX6 (32). To test if the *Hs*PEX1^G843D^ mutation impairs assembly with *Hs*PEX6 in the HCT116 cell lines, we immunoprecipitated PEX1^G843D^ from both the N- and C-terminally FLAG-tagged cell lines and identified co-immunoprecipitating proteins by mass-spectrometry. Immunoblotting of the FLAG-PEX1 co-precipitants showed a significant reduction of PEX6 co-purified with PEX1^G843D^ (Figure 5A, Figure S4A). Mass-spectrometry of both FLAG-tagged PEX1 cell lines showed a significantly reduced abundance of PEX1 binding partners PEX6 and PEX26 when G843D was present, confirming that PEX1^G843D^ impairs assembly with PEX6 (Figure 5B, Figure S4B).

**Figure 5:**
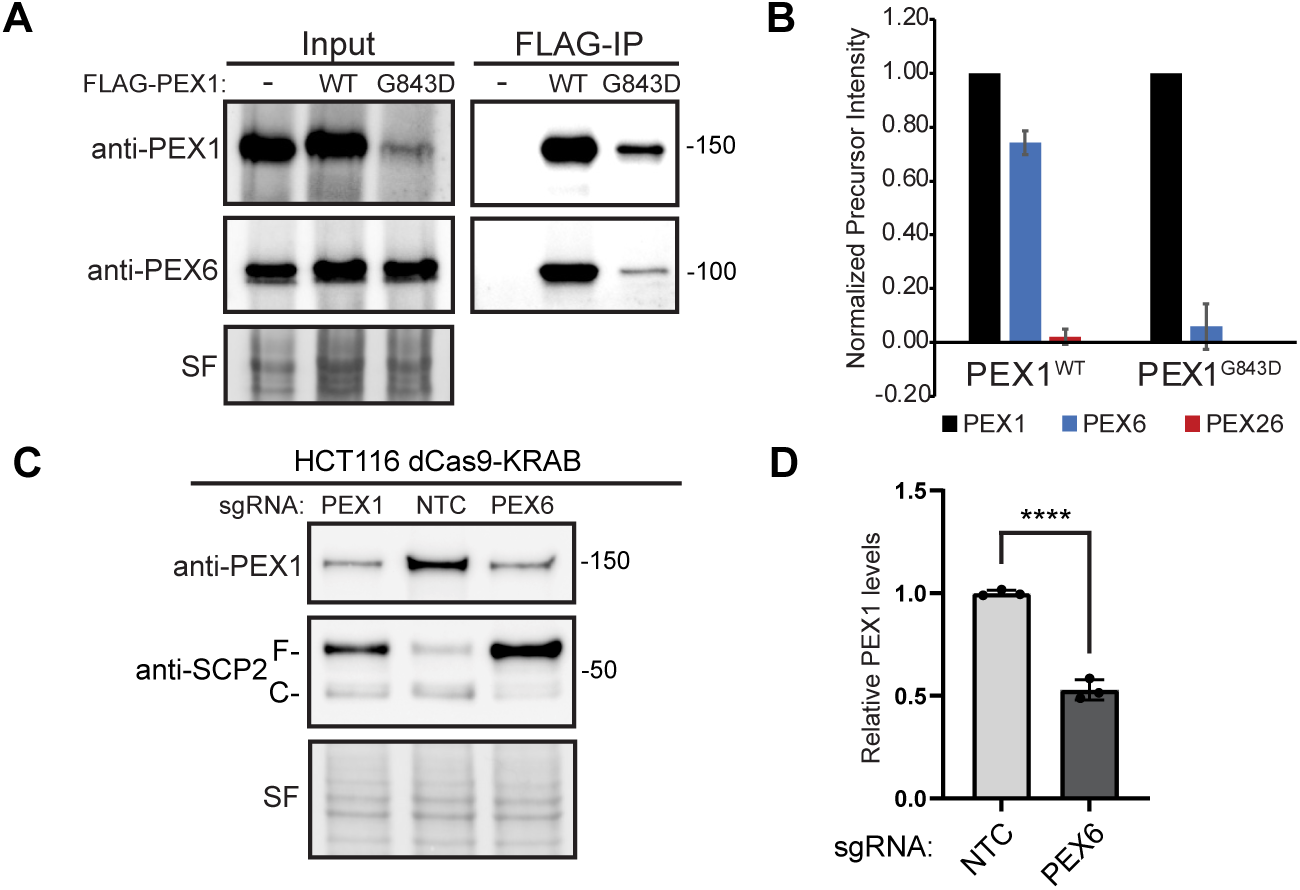
An assembly defect is sufficient to cause degradation of PEX1^WT^ and PEX1^G843D^. A) Immunoprecipitation of FLAG-PEX1 and FLAG-PEX1^G843D^ from clonal HCT116s show reduced co-immunoprecipitation of PEX6 in FLAG-PEX1^G843D^ cell lines (PEX1 = 142 kDa, PEX6 = 106 kDa, SF = StainFree total protein). B) Mass spectrometry of eluates from FLAG immunoprecipitations in FLAG-PEX1^WT^ and FLAG-PEX1^G843D^ cell lines shows a significant decrease in abundance of PEX1 binding partners PEX6 and PEX26 with the G843D mutation. Total precursor intensity was normalized to PEX1 precursor intensity levels. C) Knockdown of PEX6 with sgRNA in dCas9-KRAB expressing HCT116 cells causes degradation of PEX1^WT^ and phenocopies a PEX1 knockdown (PEX1 = 142 kDa, SF = StainFree total protein). Full-length SCP2 (SCP2-F, 58 kDa) is cleaved upon import into the peroxisome (SCP2-C, 46 kDa). E) Quantification of PEX1 levels in PEX6 knockdown and NTC cells from (C) show a significant decrease in PEX1^WT^ levels. Means with SD are shown for N=4 replicates. Unpaired t-test, p-value <0.0001.

Recent studies have demonstrated that cells degrade unpaired, or “orphan”, subunits of multi-protein assemblies in an attempt to regulate protein-complex stoichiometry (45). To test whether the assembly of the PEX1/PEX6 complex is also subject to orphan protein quality control mechanisms, we tested if depletion of PEX1’s binding partner PEX6 is sufficient to induce the degradation of PEX1^WT^. We found that knockdown of PEX6 in HCT116 CRISPRi (dCas9-KRAB)(46–48) cells significantly reduced wild-type PEX1 protein levels (Figure 5C-5D, Figure S4C-S4D). These data suggest that the PEX1/PEX6 complex is subject to a previously uncharacterized orphan protein quality control mechanism, and that wild-type PEX1 is degraded in the absence of its PEX6 binding partner. Therefore, we conclude that PEX1^G843D^ is defective in its ability to bind PEX6, which exacerbates PEX1 orphan protein quality control, likely in addition to other redundant misfolded protein quality control pathways that target PEX1^G843D^ for degradation.

### PEX1^G843D^ is stabilized by inhibiting ubiquitination

Several E3 ligases have been recently identified to monitor the formation of protein complexes and to ubiquitinate orphan proteins that have not properly assembled, such as UBE2O, HUWE1, UBR5 and SCF^Das1^ (49–52). In our co-immunoprecipitation mass-spectrometry we identified multiple E3 ligases enriched in binding to PEX1^G843D^ compared to PEX1^WT^ (Figure 6A). We generated HCT116 CRISPRi (dCas9-KRAB)(46–48) cells homozygous for the G843D mutation using the same strategy as for the FLAG-PEX1 cell lines (Figure S5A-S5B). We then knocked down two E3 ligases that were enriched in FLAG-PEX1^G843D^ eluates (Figure 6A) and have been implicated in orphan protein quality control, UBR5 and UBE2O (50, 52). We observed a modest increase in the steady state levels of PEX1 in UBR5 knockdowns, as well as a modest increase in the levels of PEX1 protein remaining 24 hours after cycloheximide treatment (Figure 6B). UBE2O knockdown appeared to have no effect on PEX1 protein levels (Figure 6B), although the knockdown of UBE2O mRNA was effective (Figure S5C). These experiments suggest that UBR5 ubiquitinates PEX1^G843D^, and that knockdown of this ligase can modestly stabilize PEX1^G843D^. We expect PEX1^G843D^ to be targeted by several E3 ligases since it has both a folding and assembly defect, so it is unsurprising that PEX1^G843D^ levels do not recover to WT levels upon knockdown of a single E3 ligase.

**Figure 6:**
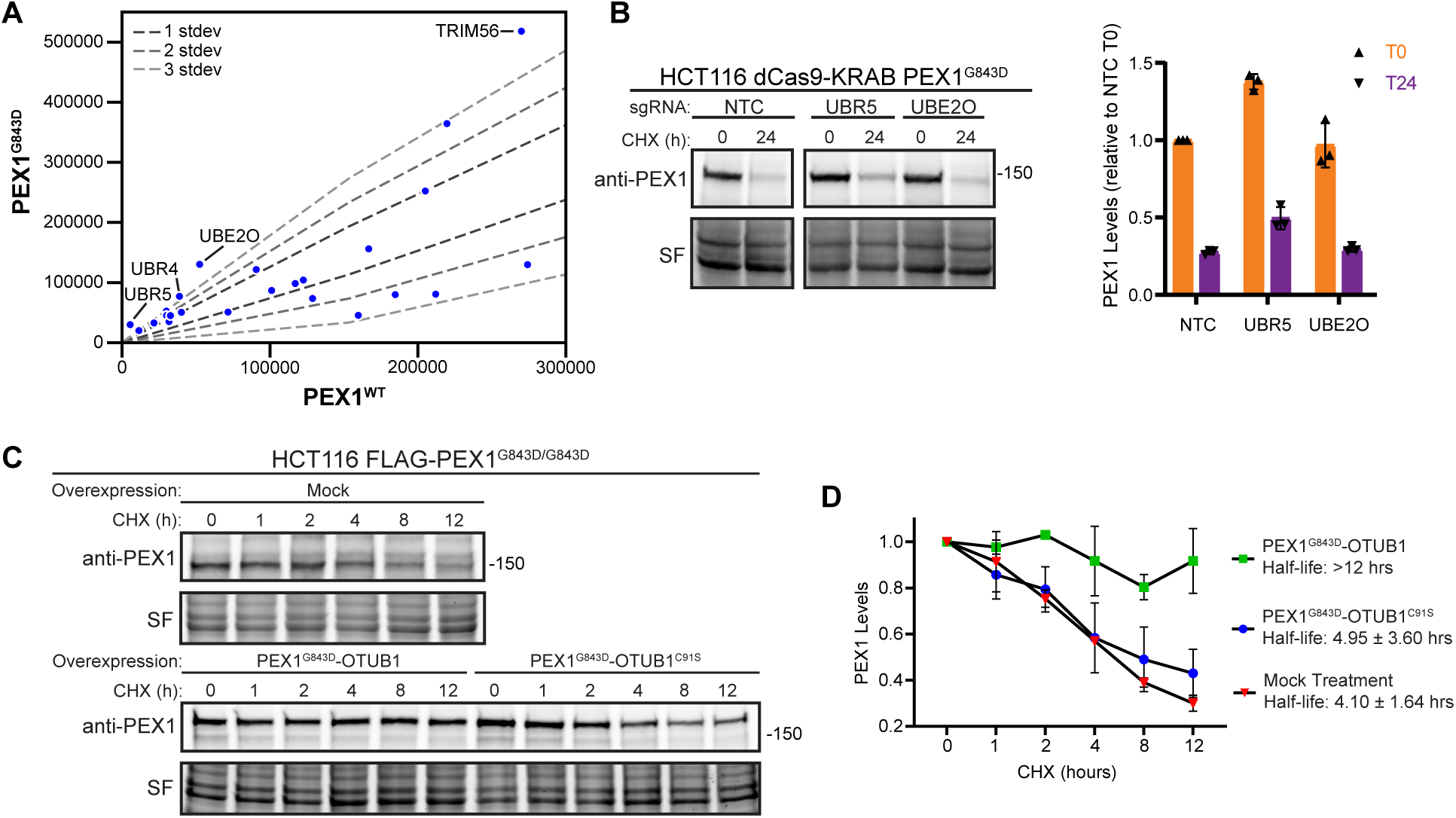
PEX1^G843D^ degradation is hindered by fusion to a deubiquitinase OTUB1. A) Ubiquitin ligase related proteins identified by anti-FLAG co-purification MS/MS in FLAG-PEX1^WT^, FLAG-PEX1^G843D^, PEX1^WT^-FLAG, and PEX1^G843D^-FLAG cell lines. The plot shows the comparison between WT and G843D samples on a subset of total identified proteins related to ubiquitin interacting proteins. B) Representative immunoblots and quantification of PEX1 levels of PEX1^G843D/G843D^ HCT116 CRISPRi (dCas9-KRAB) cell lines with knockdown of UBR5 and UBE2O candidates identified from the IP-MS experiment (PEX1 = 142 kDa, SF = StainFree total protein). Knockdown of mRNA transcripts verified in Figure S5. Cells were treated with cycloheximide at T=0 and samples collected after 24 hours. Means with SD are shown from N=3 replicates for each timepoint. C) Representative immunoblots of PEX1-OTUB1 fusions in a CHX-chase assay. OTUB1^C91S^ is catalytically inactive. D) Quantification of replicate experiments similar to (C). Means with SD are shown from N=3 replicates for each timepoint.

We therefore tested if fusion of PEX1^G843D^ to a deubiquitinase would better shield PEX1 from degradation, similar to the deubiquitinase-targeted chimera (DUBTAC) strategy that has been shown to effectively protect proteins from overzealous protein degradation (53). We generated constructs to express a chimera of PEX1^G843D^ with the deubiquitinase OTUB1 fused to the C-terminus of PEX1^G843D^. We transfected these constructs into our FLAG-PEX1^G843D^ cells and performed a cycloheximide chase assay. We found that PEX1^G843D^-OTUB1 was stable over the 12-hour course of the assay (Figure 6C). Critically, an identical chimeric construct of PEX1^G843D^ with catalytically inactive OTUB1, OTUB1^C91S^, was not stabilized in this assay (Figure 6C). Densitometry quantification of replicate immunoblot experiments show that PEX1^G843D^-OTUB1 was significantly stabilized while PEX1^G843D^-OTUB1^C91S^ was degraded at a rate similar to endogenous PEX1^G843D^ (Figure 6D). Altogether, our results suggest that degradation of PEX1^G843D^ is ubiquitin-dependent, and that targeted de-ubiquitination is sufficient to protect PEX1^G843D^ from proteasomal degradation.

## Discussion

In this study, we leveraged the tractability of the yeast Pex1/Pex6 proteins and Cas9 gene editing to explore the consequences of the G843D mutation on PEX1. Our *in vitro* biochemical analysis of the homologous yeast variant, *Sc*Pex1^G700D^, showed that this mutation directly destabilizes the D2 ATPase domain and reduces assembly with Pex6. Despite these defects, *Sc*Pex1^G700D^/Pex6 retains ATPase and unfoldase activity *in vitro*. Additionally, expression of Pex1^G700D^ mostly restored peroxisome import in the Δ*pex1* strain, suggesting that Pex1^G700D^ is functional when overexpressed. In human cells, we found that PEX1^G843D^ was subject to rapid degradation by the ubiquitin proteasome system, but when overexpressed, PEX1^G843D^ could support peroxisome function. Thus, our data show that PEX1^G843D^ retains function, but is degraded by overzealous protein quality control machinery. This work suggests that the widely varying clinical phenotypes and disease severity of patients harboring PEX1^G843D^ depends on the residual levels of PEX1 protein and is sensitive to variations in the ubiquitin-proteasome system. Future work could explore the synergistic effects of treatments that stabilize PEX1^G843D^ alongside small molecules that have been shown to improve peroxisome import in PEX1^G843D^ cell lines (40, 41, 54).

Protein quality control by ubiquitin-mediated degradation is a well-established post-translational regulatory mechanism. Recent studies have shown that cells not only monitor protein folding, but also the correct assembly of large protein complexes. Indeed, for some AAA-ATPases, such as the proteasome, assembly is a strictly regulated process with co-chaperones and assembly factors that facilitate proper pairing or degradation of each AAA-ATPase domain (55, 56). From our experiments, we surmise that PEX1^G843D^ degradation occurs in a complex manner with recognition by multiple components of the ubiquitin-proteasome system due to both intrinsic folding instability and poor complex formation with PEX6. Thus, the functional redundancy between E3 ligases precludes a strong phenotype from single knockdown, but forced proximity of a deubiquitinase clearly protected the protein from ubiquitin-proteasome degradation. Interestingly, we found that even wild-type PEX1 was subject to degradation in the absence of its binding partner PEX6. This observation suggests that assembly of PEX1 and PEX6 masks a previously unknown ‘degron’ in PEX1, and thus there may be other factors that specifically monitor PEX1/PEX6 assembly that are yet to be identified. Therefore, the degradation of orphaned PEX1 appears to be important for cellular function. Since PEX6 binds the peroxisomal anchoring protein, PEX26, orphaned PEX1 lacks the information for localization to the peroxisome and may mislocalize and interact with other Type 2 AAA-ATPases such as p97, SPATA5, and ATAD2 which were also identified as co-immunoprecipitants in our PEX1 mass spectrometry experiments. AAA-ATPases may be prone to mismatch since the conservation of the nucleotide binding sites at the interfaces of subunits likely restricts variation and specialization in these interfaces.

Understanding the proteasome-dependent regulation of the PEX1/PEX6 complex could unveil new strategies for improving clinical outcomes in patients with PEX1^G843D^ or other mutations in PEX1 that result in protein level changes and milder phenotypes. While an ideal treatment would be systemic re-introduction of PEX1^WT^ as a gene therapy, such an approach remains impractical at post-embryonic stages. Instead, strategies to stabilize the residual protein levels of PEX1 hypomorphs could be modeled on promising approaches to treating other diseases exacerbated by overzealous protein quality control such as cystic fibrosis. Key next steps in the development of these therapeutic strategies thus include the identification of small molecules that stabilize PEX1 protein folding or PEX1/PEX6 assembly; as well as discovery of small molecule ligands that bind PEX1, which can then be conjugated to small molecules that recruit endogenous deubiquitinases to the target (53). While this study used the common disease-causing allele PEX1^G843D^, many of the rare alleles in PEX1 and PEX6 also known to cause disease are associated with reduced assembly of PEX1/PEX6 and reduced levels of PEX1 or PEX6 proteins (32). Additionally, studies suggest that import into peroxisomes declines with age and that peroxisome biogenesis is modulated by viruses (57–59). Thus, strategies that improve PEX1/PEX6 function and peroxisome import may improve human health beyond PEX1^G843D^-associated Peroxisome Biogenesis Disorders.

### Experimental Procedures

#### Purification of Recombinant Proteins

Pex1-FLAG and His-Pex6 wild-type and G700D complexes were co-expressed in BL21* *E. coli* from the pETDuet and pCOLADuet vectors, respectively. His-MBP-PSP-Pex1 wild-type and G700D D2 domains were expressed in BL21* *E. coli* from the pCOLA vector. The expression strains were grown in DYT (16 g tryptone, 10 g yeast extract, and 5 g NaCl) or DYT with 0.2% dextrose (for His-MBP-PSP-Pex1 D2 domain constructs) with appropriate antibiotics at 30°C. Cultures were induced at OD_600_ = 0.6 – 0.8 with 0.3 mM IPTG before overnight incubation at 18°C or 3-hour incubation at 30°C for Pex1 D2 domain constructs. The *E. coli* were harvested at 4000g x 20 minutes at 4°C, and the pellet was resuspended in NiA buffer (30-50 mM HEPES pH 7.6, 100 mM NaCl, 100 mM KCl, 10% glycerol, 10 mM MgCl_2_, 20 mM imidazole, and 1 mM ATP (VWR #97061-226)) with lysozyme (0.2 mg/mL, MP Bio #02100831-CF) and protease inhibitors: leupeptin, pepstatin, aprotinin, and PMSF. Cells were lysed by sonication at 60% amplitude with 15 second pulses on 90 second off for a total of 180 seconds on. The total lysate was spun at 30,000g x 30 minutes at 4°C to remove insoluble cell debris, and the supernatant was transferred to conical tubes with 5 mL of pre-equilibrated Ni-NTA agarose (Thermo #88223). The cell lysate and Ni-NTA agarose were incubated with inversion at 4°C for 1 hour before the Ni-NTA agarose was batch-washed twice with 50 mL NiA buffer. The Ni-NTA agarose was resuspended in NiA and poured into gravity flow columns and washed until the flowthrough contained no protein, as judged by Bradford assay. The bound protein was then eluted with NiA containing 500 mM imidazole. For the Pex1-FLAG/His-Pex6 complexes, the Ni-NTA eluate was then added to pre-equilibrated anti-FLAG M2 agarose affinity resin (Sigma #A2220). The anti-FLAG resin was equilibrated by first washing with two sequential column volumes of GF buffer (50-60 mM HEPES pH 7.6, 50 mM NaCl, 50 mM KCl, 5 % glycerol, 10 mM MgCl_2_, and 1 mM ATP), then with three sequential column volumes of 100 mM glycine HCL pH 3.5, followed by equilibration with 5 sequential column volumes of GF. The Ni-NTA eluate was passed over the FLAG-resin twice to bind, and then was washed with GF buffer until flowthrough contained no protein, as judged by Bradford assay. The bound protein was eluted with GF and 300 μg/mL FLAG peptide (Sigma #F3290) and subsequently concentrated on a spin concentrator (30 or 100 MWCO, Millipore Sigma #UFC910008) before snap-freezing in liquid nitrogen and storing at −80°C. To separate Pex1-FLAG/His-Pex6 hexamer from other oligomers and monomers by FPLC, the concentrated FLAG eluates were loaded onto a Superose-6 size exclusion column (Cytiva #29091596) equilibrated in GF buffer. For the complex assembly analysis of wild-type and G700D complexes, equal concentrations of concentrated FLAG-elution were loaded on the FPLC and fractions of different peaks were analyzed by SDS-PAGE and Coomassie-blue staining. The concentration of protein was determined with a Bradford assay.

For His-MBP-PSP-Pex1 D2 domain constructs, the same protocol was followed through the Ni-NTA elution, where the resulting eluate was batch bound to amylose agarose resin rather than FLAG affinity resin and eluted with NiA buffer and 10 mM maltose. Eluted protein was concentrated on a spin concentrator (30 MWCO, Millipore Sigma #UFC903008). To further purify His-MBP-PSP-Pex1 wildtype and G700D D2 domains, the concentrated amylose eluates were loaded onto a Superdex 200 (Cytiva #28990944) size exclusion column in NiA buffer. Fractions of different peaks were analyzed by SDS-PAGE and Coomassie-blue staining to identify isolated His-PSP-MBP-Pex1 D2 domains. Identified fractions were subsequently concentrated on a spin concentrator (30 MWCO), snap-frozen in liquid nitrogen, and stored at −80 °C. Protein concentration was determined via Bradford assay.

To purify the cytosolic domain of *Sc*Pex15 (amino acids 1-309), we replaced the transmembrane domain and cytosolic amphipathic helix with a FLAG-6xHis and an HRV-3C PreScission protease (PSP) site between Pex15 and the affinity tags. This construct was purified with the same protocol as for Pex1/Pex6 up to the Ni-NTA elution, except without ATP in the buffers. The Ni-NTA eluate was then concentrated with spin concentrators, and HRV-3C PreScission protease (GenScript #Z03092) was added and was then dialyzed against 2 L of NiA buffer overnight at 4°C to reduce imidazole concentration. After dialysis, the eluate was then passed over fresh Ni-NTA agarose resin to orthogonally purify Pex15 with 6xHis tag removed by collecting the flowthrough.

#### Thermal Melt Denaturation

Thermal stabilities of His-MBP-PSP-Pex1 D2 wildtype and G700D domains were monitored via increasing fluorescence of SYPRO-Orange Protein Gel Stain (Sigma #200-664-3). SYPRO-Orange Protein Gel Stain is naturally quenched in solution and fluoresces upon interaction with the hydrophobic core of a protein. The reaction mixture contained 1 mg/mL of purified MBP-fusion protein, 5X SYPRO-Orange Protein Gel Stain, variable concentrations of ATP, and NiA. 0.5 M ATP was diluted in water to generate ATP solutions with the desired final concentration (0.5 mM, 1.5 mM, 2 mM, 2.5 mM, 3 mM, 5 mM, 7 mM, 8 mM, or 10 mM) in the protein reaction mixture. SYPRO-Orange fluorescence of samples prepared in a 96-well plate were read over a temperature range of 10°C to 95°C, with 0.5°C increments, in a melt curve protocol of a CFX96 Touch RT-PCR instrument (BioRad). The negative derivative of relative fluorescence units (- d(RFU)/dt) was generated in GraphPad Prism and used to create the melt curves shown. The melting temperature (Tm) was determined using the “area under the curve” analysis in GraphPad Prism to calculate the minima of each melt curve. The minimum value of the concave-up parabola corresponding to an unfolding event was determined as the melting temperature of a domain. Referenced temperature of a given domain is an average of the determined melting temperatures (N = 3) with the standard deviation shown for each. To determine the K_d_ of *Sc*PEX1^WT^ D2 domain binding to ATP, a one-site total non-linear regression was calculated in GraphPad Prism and the value shown with 95% confidence interval.

#### ATPase Activity Assay

The ATPase activity of wild-type and G700D *Sc*Pex1/*Sc*Pex6 complexes were monitored using an ATP/NADH-coupled enzymatic assay. In this assay, the regeneration of hydrolyzed ATP is coupled to the oxidation of NADH, which is measured at 340 nm (60). The reaction mixture contains 3 U/mL pyruvate kinase, 3U/mL lactate dehydrogenase, 1 mM NADH, and 7.5 mM phosphoenol pyruvate. The assays were performed in a 96-well plate and read on a SpectraMax M5 UV-Vis plate reader spectrometer over 900 seconds with 10 second time-points. For saturating conditions, we used 5 mM ATP. In each assay, the concentration of purified *Sc*Pex1/*Sc*Pex6 complexes used was to stay in the linear range throughout the duration of the experiment. For *Sc*Pex1^G700D^/*Sc*Pex6 complexes, this required higher concentrations ranging from 10-20 nM to achieve the same linearity. To determine the effect of *Sc*Pex15 inhibition, we pre-incubated *Sc*Pex1/*Sc*Pex6 complexes with *Sc*Pex15 (1-309) before addition to the ATPase assay reaction mixture.

#### Fluoroscein-5-maleimide Unfolding Assay

Substrate *Sc*Pex15 (1-309) protein was used at a final concentration of 2.5 μM, and each reaction included ATP regeneration mixture (ATP, creatine kinase, creatine phosphate). *Sc*Pex15 (1-309) was diluted using buffer1 (50 mM HEPES pH 7.5, 50 mM NaCl, 50 mM KCl, and 10 mM MgCl_2_) before the addition of Pex1/Pex6 complexes at a final concentration of 0.1-0.2 μM. The reaction mixture was incubated for 60 seconds to allow motor unfolding, then for an additional 30 seconds with 0.5 mM fluorescein-5-maleimide F5M (Fisher #50-850-980) before quenching with equal volume of quench buffer (2% SDS, 10% β-mercaptoethanol). “Urea” samples were diluted in buffer2 (50 mM HEPES pH 7.5, 50 mM NaCl, 50 mM KCl, and 10 mM MgCl_2_, and 8M urea) to a 6M urea final concentration and incubated with F5M for 10 minutes before quenching. Excess F5M was run off SDS-PAGE gels, and the gels were imaged in the fluorescein channel (Bio-Rad ChemiDoc) prior to Coomassie-blue staining.

#### Colorimetric Peroxisome Import Assay

Import assays were performed as described previously (24). In brief, a λ1*pex1* deletion strain with the violacein pathway (VioA, VioB, and VioE-SKL) genomically incorporated at the *LEU2* locus was complemented with *CEN ARS* plasmids expressing *PEX1* variants. Three colonies from each transformation were grown overnight, diluted and grown to log-phase the next day, and spotted with 5 μl at OD_600_ = 0.3 to evaluate the formation of green pigment. To quantify the fluorescence of prodeoxyviolacein metabolites, saturated cultures of yeast were lysed in acetic acid, filtered, and the fluorescence of the lysate was quantified at excitation 535 nm and emission 585 nm.

#### Cell Lines and Culture

Primary and immortalized patient fibroblasts (a gift from the laboratory of Nancy Braverman, McGill University), HCT116, HEK293T, and HCT116 dCas9-KRAB (a gift from the laboratory of J. Corn, ETH Zürich) cells were cultured in Dulbecco’s Modified Eagle Media GlutaMax (Gibco #10569010) supplemented with 10% fetal bovine serum (R&D Systems #FS11150H) and 1% penicillin/streptomycin (Gibco #15140122) and kept at 37°C and 5% CO_2_ (patient fibroblasts were cultured at 7% CO_2_) in a humidified incubator. For routine passaging, adherent cells were grown to ∼70-80% confluency, washed with DPBS (Gibco #14190-144), and subsequently treated with 0.25% trypsin-EDTA (Gibco #25-200-072) for 3-5 min in a 37°C incubator. Lifted cells were then quenched with DMEM and passaged.

HCT116 cells are human adult male colorectal carcinoma cells. HCT116 dCas9-KRAB cells were selected for uptake of sgRNA guide plasmid with 1.5 μg/ml Puromycin (Gibco # A1113803) for 1 week and were continuously cultured in 1.0 ug/mL Puromycin to maintain selection after this time.

For drug treatment conditions, cells were treated with 100 μg/mL cycloheximide (Sigma-Aldrich #C4859), 50 nM bafilomycin (Sigma-Aldrich # B1793), 10 μM carfilzomib (Selleck #PR-171) for between 1 and 24 hours. Doxycycline treatments were either 1 µg/mL or 4 µg/mL doxycycline for tetON-PEX1^WT^ and tetON-PEX1^G843D^ respectively (Sigma #D9891-5G) for 96 hours.

#### Genome Editing with CRISPR-Cas9

HCT116 cells harboring epitope FLAG-tags were generated using CRISPR-Cas9 to introduce either N-terminal or C-terminal FLAG-tags, presence of the tag was verified in bulk culture, and then cells were dilution cloned and expanded. The presence and zygosity of FLAG insertion was confirmed with junction PCR, T7E1 digest, and western blot. To introduce the G843D mutation, the FLAG-tagged HCT116 cells were edited at the PEX1 locus with CRISPR-Cas9, cells dilution cloned and expanded, and presence and zygosity of editing was confirmed with ClaI digests of PCR amplicons of the *PEX1* gene from exon 14 through exon 15 from genomic DNA. For CRISPR-Cas9 editing, cells were nucleofected by combining RNP buffer (100 mM HEPES, 750 mM KCl, 25 mM MgCl_2_, 25% glycerol, and 5 mM TCEP), 2.5 μM sgRNA, 2 μM Cas9 (purchased from Berkeley MacroLab), and 5 μM dsDNA donor, and then adding this mixture to 200k cells washed with DPBS (Gibco #14190-144) and resuspended in nucleofection buffer SE (Lonza #V4XC-1024). Reaction mixtures were then electroporated in 4D Nucleocuvettes (Lonza) and subsequently recovered with pre-warmed media in culture dishes.

#### RNA Extraction, cDNA Synthesis, and Quantitative RT-PCR

Normalized cell counts were lysed with RLT buffer with 1% 2-mercaptoethanol and RNA was extracted using the Qiagen RNEasy kit (Qiagen #74106), according to the manufacturer’s instructions. Genomic DNA in the cell lysate was pre-sheared with Qiagen QIAShredder (Qiagen #79656) columns before RNA extraction. 1 μg of RNA was reverse transcribed to generate cDNA with the iScript Reverse Transcription Supermix (BioRad #1725035) according to the manufacturers protocol. RNA was stored at −80°C and cDNA was stored at −20°C. The cDNA was diluted 10-fold in nuclease-free water and 2% was used as input for qRT-PCR using SYBR Green (BioRad #1725271) based gene-specific qPCR in a CFX96 Touch qPCR instrument (BioRad). All qPCR experiments were run with technical triplicates for each biological replicate and changes in gene expression were determined using the ΔΔCq method.

#### Immunofluorescence Staining

Cells were plated on glass bottom 96-well plates coated with either 50 μg/mL poly-D-lysine (Gibco #A3890401) or Collagen (Corning #354231) and fixed using 4% paraformaldehyde (Electron Microscopy Sciences #15710) in DPBS (Gibco #14190-144) for 10 minutes and washed with DPBS. Cells were then permeabilized using 0.1% Triton X-100 (Thermo Fisher #A16046-AP) in DPBS for 10 minutes, blocked with Buffer FZ (50 mM NH4Cl, 0.5% BSA, 0.05% saponin, 0.02% NaN3 in PBS) for 20 minutes, and then probed with desired antibody diluted 1:1000 in Buffer FZ overnight at 4C with gentle agitation. The following day, cells were washed 3X with DPBS-T and incubated with the appropriate fluorescently conjugated secondary antibody diluted 1:25000 in Buffer FZ. After secondary staining, DAPI staining (Invitrogen #D1306) was carried out for 15 minutes in DPBS. Cells were then washed 3X in DPBS and stored in DPBS prior to image acquisition. All antibodies used for immunofluorescence staining in this study are as follows: Catalase (Proteintech #66765-1-Ig), PMP70 (Invitrogen PA1-650), AlexaFluor 488 goat anti-rabbit igG [H+L] (Invitrogen #A11034), Goat anti-mouse IgG [H+L] Alexa Fluor 594 (Invitrogen #A32742).

#### Confocal Microscopy and Analysis

Confocal microscopy images were acquired using an inverted spinning disc confocal microscope (Nikon Ti-Eclipse) equipped with an electron multiplying charge-coupled device camera (Fusion SN:500241) and environmental control (Okolabs stage top incubator). Image acquisition was performed with a 100X NA 1.64 oil-immersion objective. Exposure times and laser powers were standardized between replicates of an experiment. Each experiment was analyzed in technical triplicate, as well as biological triplicate. Sixteen images were collected per technical triplicate (i.e., per well of a 96-well plate) in a fixed pattern. Raw micrographs were analyzed using the open-source image analysis software, CellProfiler. In brief, nuclei objects were identified and used to find cell borders by intensity-based edge propagation. PMP70 and catalase foci were identified, and their features quantified on a per cell basis. Pearson’s colocalization coefficient was calculated by the overlap of catalase positive pixels with PMP70 pixels and quantified on a per cell basis. All data presented contains colocalization coefficients calculated from N>200 cells. Resulting analyses were plotted using Python3.

#### Cycloheximide Chase Assay

For all CHX chase experiments, 800k – 1 million cells were seeded in 6 cm plates the day before treatment. For the autophagy inhibition experiments, cells were treated with either DMSO, 1 nM bafilomycin-A1 (Cell Signaling #54645), or 1nM bafilomycin-A1 and 100 µg/mL cycloheximide (Sigma-Aldrich #C4859) at T=0 and lysates collected at indicated time points. For the proteasome inhibition experiments, cells were pre-treated in 10 μM carfilzomib (Selleck #PR-171) for 1 hour before being released into media containing 100 μg/mL cycloheximide and 10 uM carfilzomib at T=0 and lysates were taken at each indicated time point. The Half-lives for CHX chase experiments were calculated with GraphPad Prism using one-phase decay non-linear fit with 95% CI shown.

#### Lentiviral Production and Transduction, and gene knockdown by CRISPRi

VSV-G pseudo-typed lentiviral particles were generated by transfection of HEK293T cells with second generation lentiviral packaging vectors using TransIT-LT1 Transfection Reagent (Mirus #MIR2305) according to the manufacturers protocol. Lentivirus was harvested 72h after transfection and stored at - 80°C. To perform transductions with the TetOn constructs, lentiviral supernatant was passively applied to cells overnight, followed by expansion and selection of clonal populations by FACS (Sony SH800S) for constitutive BFP expression. Clonal populations were then analyzed by flow cytometry (Attune NxT Flow Cytometer, Invitrogen) and populations expressing varying levels of BFP signal were expanded. Cells with high expression of TetOn-PEX1 were analyzed by treating cells with 2.0 μg/mL Doxycycline for 24h before analyzing expression via dot-immunoblotting with the PEX1 antibody (BD Biosciences #611719).

Knockdown of candidate genes, PEX1, and PEX6 were performed using CRISPRi in the HCT116 dCas9-KRAB cell line. Constructs were made by cloning sgRNA sequences into the backbone vector pCRISPRia-v2, a gift from Jonathan Weissman (Addgene #84832). Lentivirus containing the candidate sgRNA were generated as described above and used to transduce the HCT116 dCas9-KRAB parental or PEX1^G843D^ cell line and selected for by 1.5 μg/mL Puromycin (Gibco #A1113803) for at least one week. Gene knockdown was verified by RT-qPCR and/or immunoblotting.

#### Immunoprecipitation

Cell lines were harvested at 15-20 million cells (normalized among conditions and replicates), spun 500g x 5 minutes, washed in DPBS, and resuspended in LB1 (60 mM HEPES pH 7.6, 150 mM NaCl, 150 mM KCl, 10 mM MgCl_2_, 5% glycerol, 0.25 mM ATPγS (Biorbyt #orb64057), 1.0 % Triton X-100, and 0.5 mM EDTA) with added Benzonase and 1X Protease Inhibitor (Halt #PI78429). Cells were incubated on ice for 15 minutes, passed 5x through a 25G hypodermic needle to shear cells and genomic DNA, then incubated another 15 minutes on ice with mixing by inversion. The resulting lysate was then spun 15,000g x 30 minutes at 4°C and the supernatant was collected. Lysate was then pre-cleared with Protein G agarose beads (EMD Millipore #16-266), that were pre-washed three times in LB1, for 30 minutes at 4°C to reduce non-specific binding and the beads were spun down 5000g x 1 min and supernatant collected. Anti-FLAG M2 affinity agarose (Sigma #A2220) was washed three times in LB1, and then added to the pre-cleared lysate and incubated for 2-3 hours at 4°C. The FLAG-agarose resin was washed three times in LB1, and twice with LB1 containing no detergent. Proteins were eluted with 300 μg/mL FLAG-peptide (Sigma #F3290) dissolved in LB1 with no detergent by incubating for 30 minutes shaking at 1100 rpm at 4°C. Beads were spun 5000g x 1 min and eluates were collected. Eluates were snap-frozen in liquid nitrogen and stored at −80°C.

#### Mass Spectrometry Sample Preparation

FLAG co-immunoprecipitation eluates were prepared for mass spectrometry by first precipitating proteins with 20% TCA at −80°C overnight. The protein pellets were washed three times in ice-cold solution of 0.01 N HCl in 90% acetone and air-dried. Pellets were then resuspended in 100 mM Tris pH 8.5 and 8M urea, and 3 mM TCEP (GoldBio #51805-45-9) was added and incubated for 20 minutes at room temperature. Then, 10 mM iodoacetamide (Acros Organics #144-48-9) was added and incubated for 15 minutes at room temperature in the dark. Samples were then diluted 4-fold with 100 mM Tris pH 8.5, and 1 mM CaCl_2_ was added along with 1 μg of Trypsin (Promega #V5280) and incubated overnight at 37°C. Digested protein samples were then snap-frozen in liquid nitrogen and sent to UC Davis Proteomics Core Facility for analysis.

#### Mass Spectrometry (LC-MS/MS)

Liquid chromatography was performed on an ultra-high pressure liquid chromatography system, nanoElute (Bruker Daltonics), at 40 °C and with a constant flow of 400 nL/min on a PepSep 150 μm x 25 cm C18 column (PepSep, Denmark) with 1.5 μm particle size (100 Å pores) (Bruker Daltonics), and a ZDV spray emitter (Bruker Daltonics). Mobile phases A and B were water with 0.1% formic acid (v/v) and 80/20/0.1% acetonitrile/water/formic acid (v/v/v), respectively. Peptides were separated using a 60 min gradient.

Mass Spectrometry was performed on a hybrid trapped ion mobility spectrometry-quadrupole time of flight mass spectrometer (timsTOF pro, Bruker Daltonics) with a modified nano-electrospray ion source (CaptiveSpray, Bruker Daltonics). The mass spectrometer was operated in PASEF mode. Desolvated ions entered the vacuum region through the glass capillary and deflected into the TIMS tunnel which is electrically separated into two parts (dual TIMS). The first region is operated as an ion accumulation trap that primarily stores all ions entering the mass spectrometer, while the second part performs trapped ion mobility analysis.

A single TIMS-MS scan is composed of many individual TOF scans of about 110 s each. All experiments were acquired with a 100 ms ramp and 10 PASEF MS/MS scans per topN acquisition cycle. In TOF mass spectrometry, signal-to-noise ratios can conveniently be increased by summation of individual TOF scans. Here, low-abundance precursors with an intensity below a ‘target value’ were repeatedly scheduled for PASEF-MS/MS scans until the summed ion count reached the target value (e.g. four times for a precursor with the intensity 5000 arbitrary units (a.u.) and a target value of 20,000 a.u. We set the target value to 20,000 a.u. for all methods. MS and MS/MS spectra were recorded from m/z 100 to 1700. A polygon filter was applied to the m/z and ion mobility plane to select features most likely representing peptide precursors rather than singly charged background ions. The quadrupole isolation width was set to 2 Th for m/z under 700 and 3 Th for m/z larger than 700, and the collision energy was ramped stepwise as a function of increasing ion mobility: 52 eV for 0–19% of the ramp time; 47 eV from 19–38%; 42 eV from 38–57%; 37 eV from 57–76%; and 32 eV for the remainder.

#### Mass Spectrometry Data Analysis

All MS/MS samples were analyzed using MSFragger (61). MSFragger was set up to search the 2021-11-16-decoys-contam-UP000005640 database assuming the digestion enzyme stricttrypsin. MSFragger was searched with a fragment ion mass tolerance of 20 PPM and a parent ion tolerance of 20 PPM. Carbamidomethyl of cysteine was specified in MSFragger as a fixed modification. Oxidation of methionine and acetyl of the n-terminus were specified in MSFragger as variable modifications.

Criteria for protein identification: Scaffold (version Scaffold_5.0.1, Proteome Software Inc., Portland, OR) was used to validate MS/MS based peptide and protein identifications. Peptide identifications were accepted if they could be established at greater than 0.0% probability by the Peptide Prophet algorithm (62). Protein identifications were accepted if they could be established at greater than 20.0% probability and contained at least 2 identified peptides. Protein probabilities were assigned by the Protein Prophet algorithm (63). Proteins that contained similar peptides and could not be differentiated based on MS/MS analysis alone were grouped to satisfy the principles of parsimony. Proteins sharing significant peptide evidence were grouped into clusters.

#### Transient transfection

Transient transfection plasmids containing the PEX1^G843D^-OTUB1 fusions were cloned by Gibson assembly to insert PEX1^G843D^ into the transient transfection OTUB1 or OTUB1^C91S^ pcDNA3.1 plasmids (Addgene #118209 and Addgene #118210, respectively). FLAG-PEX1^G843D/G843D^ cells were transfected with the PEX1^G843D^-OTUB1 fusion plasmids using Mirus TransIT-LT1 (Mirus #MIR2305) in 10 cm plates 48h prior to treatment with CHX. 24h prior to CHX treatment, transfected cells were lifted and seeded into 6 cm plates for each timepoint. Cells were then treated with CHX at T0 and harvested at the indicated time points.

#### Immunoblotting

Normalized cell counts were lysed in either (1) RIPA buffer containing benzonase (Millipore Sigma #70664) and protease inhibitor cocktail (Halt #PI78429) for 30 min on ice before combining with 2X Laemmli sample buffer containing 2-mercaptoethanol or (2) directly in 2X Laemmli sample buffer containing 2-mercaptoethanol before sonicating for 30sec at 50% amplitude (Branson) and boiling for 5 minutes at 95°C. For the CHX chase experiments, cells were lysed with 2X Laemmli sample buffer with 2-mercaptoethanol directly in culture plates and scraped to collect cells. Samples were resolved with either 4-20% SDS-PAGE gels (Bio-Rad #4561095) or homemade bis-Tris MES-Tris Gels. Proteins were transferred to 0.45 µm LF PVDF membranes using semi-dry (BioRad) or wet-transfer methods with Towbin transfer buffer (192 mM Glycine, 25 mM Tris-base, 20% methanol). Membranes were blocked in 3% BSA (Fisher Scientific #BP1605-100) in TBST (0.1% Tween-20) for 1 hour, then probed with desired primary antibody in 3% BSA in TBST overnight at 4°C with gentle agitation. The next day membranes were washed 3×5 min each with TBST before probing with species-specific HRP-conjugated secondary antibodies dissolved in 5% milk or 3% BSA in TBST for 1 hour. Membranes were washed 3×5 min each with TBST before visualization of chemiluminescence using either Pierce ECL2 Western Blotting Substrate (Thermo Scientific #PI80196) or Pierce SuperSignal West Femto Substrate (Thermo Scientific #34095) on a ChemiDoc MP imaging system (BioRad). The antibodies used for immunoblotting in this study are as follows: PEX1 (BD Biosciences #61171), FLAG M2 (Sigma Aldrich #F1804), LC3I-II (Abcam #ab192890), Hif1α (BD Biosciences #610958), SCP-2 (Sigma-Aldrich #HPA027317), PEX6 (Santa Cruz Biotechnology #sc-271813).

#### Densitometry

Western Blot images were quantified by pixel densitometry using FiJi. All analyses were performed on raw unadjusted images. All analyses presented are normalized to the total pixel density in the lane observed in the Stain-Free blot image, representing total protein load in each lane. Stain-Free gel technology provides a greater linearity and wider dynamic range for total protein quantifications as a loading control compared to housekeeping gene controls.

#### Statistical Analyses

All statistical analyses were performed using GraphPad Prism and descriptive statistics are described in figure legends where relevant.

## Data availability

## Supporting information

## Acknowledgements

We thank Dr. Nancy Braverman and Dr. Catherine Argyriou for the gift of PBD patient fibroblast cell lines. We thank Dr. Meghan Morrissey for allowing us to utilize the lab’s spinning disc confocal microscope that was essential to this study. We thank the UC Davis Proteomics Core, supported by NIH S10 Grant S10OD026918-01A1, for their assistance with mass spectrometry analysis. Brooke Gardner acknowledges support from K99/R00GM121880, R35GM146784, and the Searle Scholars Program. Connor Sheedy acknowledges support from the UC Santa Barbara Chancellor’s Fellowship. Soham Chowdhury acknowledges support from the Storke Family Fellowship. Chris Richardson acknowledges support from R35 GM142975. The content is solely the responsibility of the authors and does not necessarily represent the official views of the National Institutes of Health. We thank the members of the Gardner lab, as well as the members of the Richardson lab, particularly Christine Joyce, for insightful discussions and comments.

## Author contributions

C.J. Sheedy: Conceptualization, Methodology, Validation, Formal analysis, Investigation, Writing – Original draft, Writing – Review and editing, Visualization.

S.P. Chowdhury: Conceptualization, Methodology, Validation, Formal analysis, Investigation, Writing – Review and editing

B.A. Ali: Investigation, Validation

J. Miyamoto: Investigation, Validation

E. Pang: Investigation, Validation

J. Bacal: Investigation, Validation

K.U. Tavasoli: Investigation

C.D. Richardson: Conceptualization, Methodology, Investigation, Supervision

B.M. Gardner: Conceptualization, Funding acquisition, Project administration, Supervision, Visualization, Investigation, Writing – original draft, Writing – review & editing.

## References

1. Jansen, R. L. M., Santana-Molina, C., van den Noort, M., Devos, D. P., and van der Klei, I. J. (2021) Comparative Genomics of Peroxisome Biogenesis Proteins: Making Sense of the PEX Proteins. Front. Cell Dev. Biol. 10.3389/fcell.2021.654163

2. Braverman, N. E., and Moser, A. B. (2012) Functions of plasmalogen lipids in health and disease. Biochimica et Biophysica Acta (BBA) - Molecular Basis of Disease. 1822, 1442–1452

3. Gabaldón, T. (2010) Peroxisome diversity and evolution. Philosophical Transactions of the Royal Society B: Biological Sciences. 365, 765–773

4. Islinger, M., Cardoso, M. J. R., and Schrader, M. (2010) Be different—The diversity of peroxisomes in the animal kingdom. Biochimica et Biophysica Acta (BBA) - Molecular Cell Research. 1803, 881– 897

5. Nordgren, M., and Fransen, M. (2014) Peroxisomal metabolism and oxidative stress. Biochimie. 98, 56–62

6. Schrader, M., Kamoshita, M., and Islinger, M. (2020) Organelle interplay—peroxisome interactions in health and disease. Journal of Inherited Metabolic Disease. 43, 71–89

7. Di Cara, F., Sheshachalam, A., Braverman, N. E., Rachubinski, R. A., and Simmonds, A. J. (2017) Peroxisome-Mediated Metabolism Is Required for Immune Response to Microbial Infection. Immunity. 47, 93–106.e7

8. Dixit, E., Boulant, S., Zhang, Y., Lee, A. S. Y., Odendall, C., Shum, B., Hacohen, N., Chen, Z. J., Whelan, S. P., Fransen, M., Nibert, M. L., Superti-Furga, G., and Kagan, J. C. (2010) Peroxisomes Are Signaling Platforms for Antiviral Innate Immunity. Cell. 141, 668–681

9. Wanders, R. J. A., Baes, M., Ribeiro, D., Ferdinandusse, S., and Waterham, H. R. (2022) The physiological functions of human peroxisomes. Physiological Reviews. 10.1152/physrev.00051.2021

10. Berger, J., Dorninger, F., Forss-Petter, S., and Kunze, M. (2016) Peroxisomes in brain development and function. Biochimica et Biophysica Acta (BBA) - Molecular Cell Research. 1863, 934–955

11. Jo, D. S., Park, N. Y., and Cho, D.-H. (2020) Peroxisome quality control and dysregulated lipid metabolism in neurodegenerative diseases. Exp Mol Med. 52, 1486–1495

12. Smith, J. J., and Aitchison, J. D. (2013) Peroxisomes take shape. Nature Reviews Molecular Cell Biology. 14, 803–817

13. Waterham, H. R., and Ebberink, M. S. (2012) Genetics and molecular basis of human peroxisome biogenesis disorders. Biochimica et Biophysica Acta (BBA) - Molecular Basis of Disease. 1822, 1430–1441

14. Braverman, N. E., Raymond, G. V., Rizzo, W. B., Moser, A. B., Wilkinson, M. E., Stone, E. M., Steinberg, S. J., Wangler, M. F., Rush, E. T., Hacia, J. G., and Bose, M. (2016) Peroxisome biogenesis disorders in the Zellweger spectrum: An overview of current diagnosis, clinical manifestations, and treatment guidelines. Molecular Genetics and Metabolism. 117, 313–321

15. Steinberg, S. J., Dodt, G., Raymond, G. V., Braverman, N. E., Moser, A. B., and Moser, H. W. (2006) Peroxisome biogenesis disorders. Biochimica et Biophysica Acta (BBA) - Molecular Cell Research. 1763, 1733–1748

16. Raas-Rothschild, A., Wanders, R. J. A., Mooijer, P. A. W., Gootjes, J., Waterham, H. R., Gutman, A., Suzuki, Y., Shimozawa, N., Kondo, N., Eshel, G., Espeel, M., Roels, F., and Korman, S. H. (2002) A PEX6-Defective Peroxisomal Biogenesis Disorder with Severe Phenotype in an Infant, versus Mild Phenotype Resembling Usher Syndrome in the Affected Parents. American Journal of Human Genetics. 70, 1062

17. Saffian, D., Grimm, I., Girzalsky, W., and Erdmann, R. (2012) ATP-dependent assembly of the heteromeric Pex1p–Pex6p-complex of the peroxisomal matrix protein import machinery. Journal of Structural Biology. 179, 126–132

18. Tamura, S., Yasutake, S., Matsumoto, N., and Fujiki, Y. (2006) Dynamic and Functional Assembly of the AAA Peroxins, Pex1p and Pex6p, and Their Membrane Receptor Pex26p*. Journal of Biological Chemistry. 281, 27693–27704

19. Meyer, H., Bug, M., and Bremer, S. (2012) Emerging functions of the VCP/p97 AAA-ATPase in the ubiquitin system. Nat Cell Biol. 14, 117–123

20. Zhao, C., Slevin, J. T., and Whiteheart, S. W. (2007) Cellular functions of NSF: Not just SNAPs and SNAREs. FEBS Letters. 581, 2140–2149

21. Pedrosa, A. G., Francisco, T., Bicho, D., Dias, A. F., Barros-Barbosa, A., Hagmann, V., Dodt, G., Rodrigues, T. A., and Azevedo, J. E. (2018) Peroxisomal monoubiquitinated PEX5 interacts with the AAA ATPases PEX1 and PEX6 and is unfolded during its dislocation into the cytosol. Journal of Biological Chemistry. 293, 11553–11563

22. Platta, H. W., Grunau, S., Rosenkranz, K., Girzalsky, W., and Erdmann, R. (2005) Functional role of the AAA peroxins in dislocation of the cycling PTS1 receptor back to the cytosol. Nat Cell Biol. 7, 817–822

23. Romano, F. B., Blok, N. B., and Rapoport, T. A. (2019) Peroxisome protein import recapitulated in Xenopus egg extracts. Journal of Cell Biology. 218, 2021–2034

24. Gardner, B. M., Castanzo, D. T., Chowdhury, S., Stjepanovic, G., Stefely, M. S., Hurley, J. H., Lander, G. C., and Martin, A. (2018) The peroxisomal AAA-ATPase Pex1/Pex6 unfolds substrates by processive threading. Nat Commun. 9, 135

25. Knoops, K., de Boer, R., Kram, A., and van der Klei, I. J. (2015) Yeast pex1 cells contain peroxisomal ghosts that import matrix proteins upon reintroduction of Pex1. J Cell Biol. 211, 955– 962

26. Law, K. B., Bronte-Tinkew, D., Di Pietro, E., Snowden, A., Jones, R. O., Moser, A., Brumell, J. H., Braverman, N., and Kim, P. K. (2017) The peroxisomal AAA ATPase complex prevents pexophagy and development of peroxisome biogenesis disorders. Autophagy. 13, 868–884

27. Yu, H., Kamber, R. A., and Denic, V. (2022) The peroxisomal exportomer directly inhibits phosphoactivation of the pexophagy receptor Atg36 to suppress pexophagy in yeast. eLife. 11, e74531

28. Ebberink, M. S., Mooijer, P. A. W., Gootjes, J., Koster, J., Wanders, R. J. A., and Waterham, H. R. (2011) Genetic classification and mutational spectrum of more than 600 patients with a Zellweger syndrome spectrum disorder. Human Mutation. 32, 59–69

29. Walter, C., Gootjes, J., Mooijer, P. A., Portsteffen, H., Klein, C., Waterham, H. R., Barth, P. G., Epplen, J. T., Kunau, W.-H., Wanders, R. J. A., and Dodt, G. (2001) Disorders of Peroxisome Biogenesis Due to Mutations in *PEX1*: Phenotypes and PEX1 Protein Levels. The American Journal of Human Genetics. 69, 35–48

30. Imamura, A., Tamura, S., Shimozawa, N., Suzuki, Y., Zhang, Z., Tsukamoto, T., Orii, T., Kondo, N., Osumi, T., and Fujiki, Y. (1998) Temperature-Sensitive Mutation in PEX1 Moderates the Phenotypes of Peroxisome Deficiency Disorders. Human Molecular Genetics. 7, 2089–2094

31. Reuber, B. E., Germain-Lee, E., Collins, C. S., Morrell, J. C., Ameritunga, R., Moser, H. W., Valle, D., and Gould, S. J. (1997) Mutations in PEX1 are the most common cause of peroxisome biogenesis disorders. Nat Genet. 17, 445–448

32. Geisbrecht, B. V., Collins, C. S., Reuber, B. E., and Gould, S. J. (1998) Disruption of a PEX1–PEX6 interaction is the most common cause of the neurologic disorders Zellweger syndrome, neonatal adrenoleukodystrophy, and infantile Refsum disease. Proceedings of the National Academy of Sciences. 95, 8630–8635

33. Maxwell, M. A., Allen, T., Solly, P. B., Svingen, T., Paton, B. C., and Crane, D. I. (2002) Novel PEX1 mutations and genotype–phenotype correlations in Australasian peroxisome biogenesis disorder patients. Human Mutation. 20, 342–351

34. Klouwer, F. C. C., Falkenberg, K. D., Ofman, R., Koster, J., van Gent, D., Ferdinandusse, S., Wanders, R. J. A., and Waterham, H. R. (2021) Autophagy Inhibitors Do Not Restore Peroxisomal Functions in Cells With the Most Common Peroxisome Biogenesis Defect. Front. Cell Dev. Biol. 10.3389/fcell.2021.661298

35. Ali, B. A., Judy, R. M., Chowdhury, S., Jacobsen, N. K., Castanzo, D. T., Carr, K. L., Richardson, C. D., Lander, G. C., Martin, A., and Gardner, B. M. (2024) The N1 domain of the peroxisomal AAA-ATPase Pex6 is required for Pex15 binding and proper assembly with Pex1. Journal of Biological Chemistry. 300, 105504

36. Rüttermann, M., Koci, M., Lill, P., Geladas, E. D., Kaschani, F., Klink, B. U., Erdmann, R., and Gatsogiannis, C. (2023) Structure of the peroxisomal Pex1/Pex6 ATPase complex bound to a substrate. Nat Commun. 14, 5942

37. Jumper, J., Evans, R., Pritzel, A., Green, T., Figurnov, M., Ronneberger, O., Tunyasuvunakool, K., Bates, R., Žídek, A., Potapenko, A., Bridgland, A., Meyer, C., Kohl, S. A. A., Ballard, A. J., Cowie, A., Romera-Paredes, B., Nikolov, S., Jain, R., Adler, J., Back, T., Petersen, S., Reiman, D., Clancy, E., Zielinski, M., Steinegger, M., Pacholska, M., Berghammer, T., Bodenstein, S., Silver, D., Vinyals, O., Senior, A. W., Kavukcuoglu, K., Kohli, P., and Hassabis, D. (2021) Highly accurate protein structure prediction with AlphaFold. Nature. 10.1038/s41586-021-03819-2

38. Judy, R. M., Sheedy, C. J., and Gardner, B. M. (2022) Insights into the Structure and Function of the Pex1/Pex6 AAA-ATPase in Peroxisome Homeostasis. Cells. 11, 2067

39. Schieferdecker, A., and Wendler, P. (2019) Structural Mapping of Missense Mutations in the Pex1/Pex6 Complex. International Journal of Molecular Sciences. 20, 3756

40. MacLean, G. E., Argyriou, C., Di Pietro, E., Sun, X., Birjandian, S., Saberian, P., Hacia, J. G., and Braverman, N. E. (2019) Zellweger spectrum disorder patient–derived fibroblasts with the PEX1-Gly843Asp allele recover peroxisome functions in response to flavonoids. Journal of Cellular Biochemistry. 120, 3243–3258

41. Zhang, R., Chen, L., Jiralerspong, S., Snowden, A., Steinberg, S., and Braverman, N. (2010) Recovery of PEX1-Gly843Asp peroxisome dysfunction by small-molecule compounds. Proceedings of the National Academy of Sciences. 107, 5569–5574

42. DeLoache, W. C., Russ, Z. N., and Dueber, J. E. (2016) Towards repurposing the yeast peroxisome for compartmentalizing heterologous metabolic pathways. Nat Commun. 7, 11152

43. Preuss, N., Brosius, U., Biermanns, M., Muntau, A. C., Conzelmann, E., and Gärtner, J. (2002) PEX1 Mutations in Complementation Group 1 of Zellweger Spectrum Patients Correlate with Severity of Disease. Pediatr Res. 51, 706–714

44. Stolowich, N. J., Petrescu, A. D., Huang, H., Martin, G. G., Scott, A. I., and Schroeder, F. (2002) Sterol carrier protein-2: structure reveals function. Cell Mol Life Sci. 59, 193–212

45. Juszkiewicz, S., and Hegde, R. S. (2018) Quality Control of Orphaned Proteins. Molecular Cell. 71, 443–457

46. Gilbert, L. A., Horlbeck, M. A., Adamson, B., Villalta, J. E., Chen, Y., Whitehead, E. H., Guimaraes, C., Panning, B., Ploegh, H. L., Bassik, M. C., Qi, L. S., Kampmann, M., and Weissman, J. S. (2014) Genome-Scale CRISPR-Mediated Control of Gene Repression and Activation. Cell. 159, 647–661

47. Liang, J. R., Lingeman, E., Ahmed, S., and Corn, J. E. (2018) Atlastins remodel the endoplasmic reticulum for selective autophagy. Journal of Cell Biology. 217, 3354–3367

48. Vu, J. T., Tavasoli, K. U., Sheedy, C. J., Chowdhury, S. P., Mandjikian, L., Bacal, J., Morrissey, M. A., Richardson, C. D., and Gardner, B. M. (2024) A genome-wide screen links peroxisome regulation with Wnt signaling through RNF146 and TNKS/2. Journal of Cell Biology. 223, e202312069

49. Kong, K.-Y. E., Shankar, S., Rühle, F., and Khmelinskii, A. (2023) Orphan quality control by an SCF ubiquitin ligase directed to pervasive C-degrons. Nat Commun. 14, 8363

50. Mark, K. G., Kolla, S., Garshott, D. M., Martínez-González, B., Xu, C., Akopian, D., Haakonsen, D. L., See, S. K., and Rapé, M. (2022) Orphan quality control shapes network dynamics and gene expression. 10.1101/2022.11.06.515368

51. Xu, Y., Anderson, D. E., and Ye, Y. (2016) The HECT domain ubiquitin ligase HUWE1 targets unassembled soluble proteins for degradation. Cell Discov. 2, 16040

52. Yanagitani, K., Juszkiewicz, S., and Hegde, R. S. (2017) UBE2O is a quality control factor for orphans of multiprotein complexes. Science. 357, 472–475

53. Henning, N. J., Boike, L., Spradlin, J. N., Ward, C. C., Liu, G., Zhang, E., Belcher, B. P., Brittain, S. M., Hesse, M. J., Dovala, D., McGregor, L. M., Valdez Misiolek, R., Plasschaert, L. W., Rowlands, D. J., Wang, F., Frank, A. O., Fuller, D., Estes, A. R., Randal, K. L., Panidapu, A., McKenna, J. M., Tallarico, J. A., Schirle, M., and Nomura, D. K. (2022) Deubiquitinase-targeting chimeras for targeted protein stabilization. Nat Chem Biol. 18, 412–421

54. Berendse, K., Ebberink, M. S., IJlst, L., Poll-The, B. T., Wanders, R. A., and Waterham, H. R. (2013) Arginine improves peroxisome functioning in cells from patients with a mild peroxisome biogenesis disorder. Orphanet J Rare Dis. 8, 138

55. Funakoshi, M., Robert J Tomko, J., Kobayashi, H., and Hochstrasser, M. (2009) Multiple Assembly Chaperones Govern Biogenesis of the Proteasome Regulatory Particle Base. Cell. 137, 887

56. Kaneko, T., Hamazaki, J., Iemura, S., Sasaki, K., Furuyama, K., Natsume, T., Tanaka, K., and Murata, S. (2009) Assembly Pathway of the Mammalian Proteasome Base Subcomplex Is Mediated by Multiple Specific Chaperones. Cell. 137, 914–925

57. Beltran, P. M. J., Cook, K. C., Hashimoto, Y., Galitzine, C., Murray, L. A., Vitek, O., and Cristea, I. M. (2018) Infection-induced peroxisome biogenesis is a metabolic strategy for herpesvirus replication. Cell host & microbe. 24, 526

58. Legakis, J. E., Koepke, J. I., Jedeszko, C., Barlaskar, F., Terlecky, L. J., Edwards, H. J., Walton, P. A., and Terlecky, S. R. (2002) Peroxisome Senescence in Human Fibroblasts. MBoC. 13, 4243– 4255

59. Narayan, V., Ly, T., Pourkarimi, E., Murillo, A. B., Gartner, A., Lamond, A. I., and Kenyon, C. (2016) Deep Proteome Analysis Identifies Age-Related Processes in C. elegans. Cell Systems. 3, 144–159

60. Nørby, J. G. (1988) Coupled assay of Na+,K+-ATPase activity. in Methods in Enzymology, pp. 116–119, Biomembranes Part P: ATP-Driven Pumps and Related Transport: The Na,K-Pump, Academic Press, 156, 116–119

61. Kong, A. T., Leprevost, F. V., Avtonomov, D. M., Mellacheruvu, D., and Nesvizhskii, A. I. (2017) MSFragger: ultrafast and comprehensive peptide identification in mass spectrometry–based proteomics. Nat Methods. 14, 513–520

62. Keller, A., Nesvizhskii, A. I., Kolker, E., and Aebersold, R. (2002) Empirical Statistical Model To Estimate the Accuracy of Peptide Identifications Made by MS/MS and Database Search. Anal. Chem. 74, 5383–5392

63. Nesvizhskii, A. I., Keller, A., Kolker, E., and Aebersold, R. (2003) A Statistical Model for Identifying Proteins by Tandem Mass Spectrometry. Anal. Chem. 75, 4646–4658

64. Pan, M., Yu, Y., Ai, H., Zheng, Q., Xie, Y., Liu, L., and Zhao, M. (2021) Mechanistic insight into substrate processing and allosteric inhibition of human p97. Nat Struct Mol Biol. 28, 614–625

