## Supplementary Figures for "PEX1^G843D^ remains functional in peroxisome biogenesis but is rapidly degraded by the proteasome"

***Figure S1: Confidence metrics for AlphaFold2 predictions.***
Full length single chain predictions were obtained from the AlphaFold2 database, and the AF-ID is indicated for each. The metrics shown include the predicted aligned error maps as well as the structures and the confidence of predictions (pLDDT) color mapped onto each structure.

### Figure S1

| | | Predicted Aligned Error | 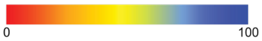  Per-Residue pLDDT |
|---|---|---|---|
| *Sc*Pex1 | Figure 1A, 1B, 1C<br>mean pLDDT: 74<br>AF-P24004 | 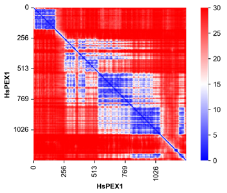  | 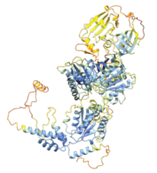 |
| *Hs*PEX1 | Figure 1B, 1C<br>mean pLDDT: 67<br>AF-O43933 | 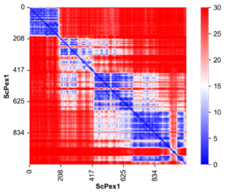  | 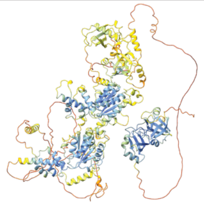 |

***Figure S2: Purification and ATPase activity of recombinant ScPex1/ScPex6 complexes.***
A) Representative two-step purification of *Sc*Pex1-FLAG/6xHis-*Sc*Pex6 and *Sc*Pex1$^{G700D}$-FLAG/6xHis-*Sc*Pex6. Pex1 and Pex6 were co-expressed in *E. coli* and purified with Ni-NTA agarose followed by anti-FLAG affinity agarose. TOT = Total lysate supernatant, FT = Flowthrough, EL = Elution. B) Representative NADH-ATP-coupled enzymatic assays for purified *Sc*Pex1/*Sc*Pex6 complexes shown in (A). The regeneration of hydrolyzed ATP is coupled to the oxidation of NADH which is read out at 340 nm.

### Figure S2

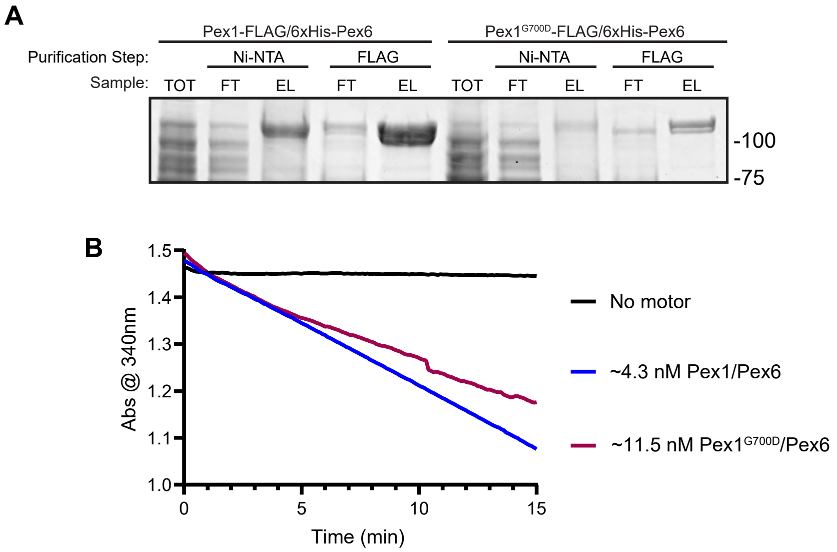

***Figure S3: Primary patient fibroblasts and strategy for CRISPR-Cas9 gene editing.***
A) Representative immunoblot of PEX1 protein levels in primary patient-derived fibroblasts with the genotypes $PEX1^{+/+}$, $PEX1^{+/G843D}$, $PEX1^{G843D/G843D}$, and $PEX1^{-/-}$. B) Strategy for CRISPR-Cas9 gene editing and selection to endogenously insert the G843D mutation into HCT116 cells. The homology donor template used for genome editing contains two base changes, one for the G843D mutation, and another silent mutation for the insertion of a ClaI restriction enzyme cut site used for screening clones. C) Representative ClaI digest of CRISPR-Cas9 edited clones to test for zygosity of G843D insertion. A 526 bp region around G843D allele was amplified by PCR from genomic DNA, followed by ClaI digestion. Full cleavage of the PCR product indicates homozygous incorporation of G843D allele.

### Figure S3

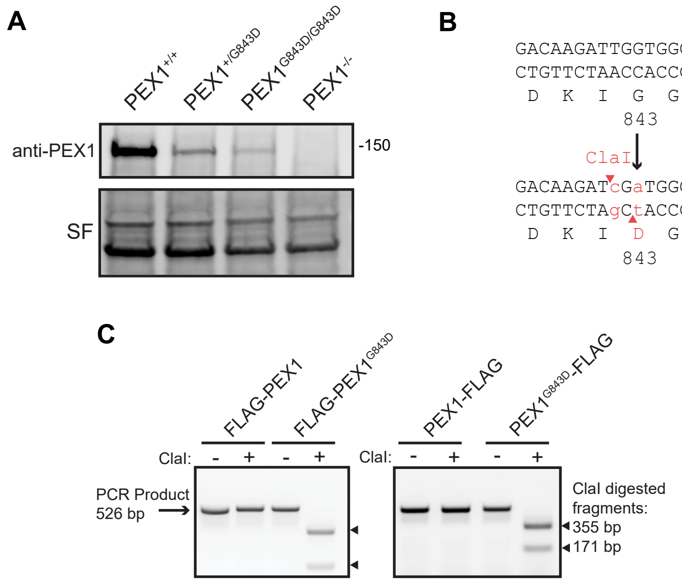

***Figure S4: Co-immunoprecipitation mass spectrometry and validation of PEX1 and PEX6 CRISPRi knockdown cells.***

A) Representative immunoblots of a co-immunoprecipitation of PEX1 from FLAG-PEX1 and PEX1-FLAG cell lines (PEX1 = 142 kDa, SF = StainFree total protein). B) Mass spectrometry on the FLAG-coimmunoprecipitations in PEX1-FLAG and PEX1$^{G843D}$-FLAG cell lines shows a decrease in abundance of PEX1 binding partners PEX6 and PEX26 with the G843D mutation. Total precursor intensity was normalized to PEX1 precursor intensity levels. C) RT-qPCR of cell lines with knockdown of either PEX1 or PEX6. CRISPRi (dCas9-KRAB) expressing cells were transduced with plasmids containing sgRNA for either PEX1 or PEX6. Means with SD are shown for N≥3 replicates. Unpaired t-test with Welch's correction. *PEX1* mRNA p-value summary: ** = 0.0044. PEX6 mRNA p-value summary: **** < 0.0001. D) Representative immunoblots of the PEX6 CRISPRi knockdown cell lines shows lower levels of PEX1 protein when PEX6 is absent (PEX1 = 142 kDa, PEX6 = 106 kDa, SF = StainFree total protein).

### Figure S4

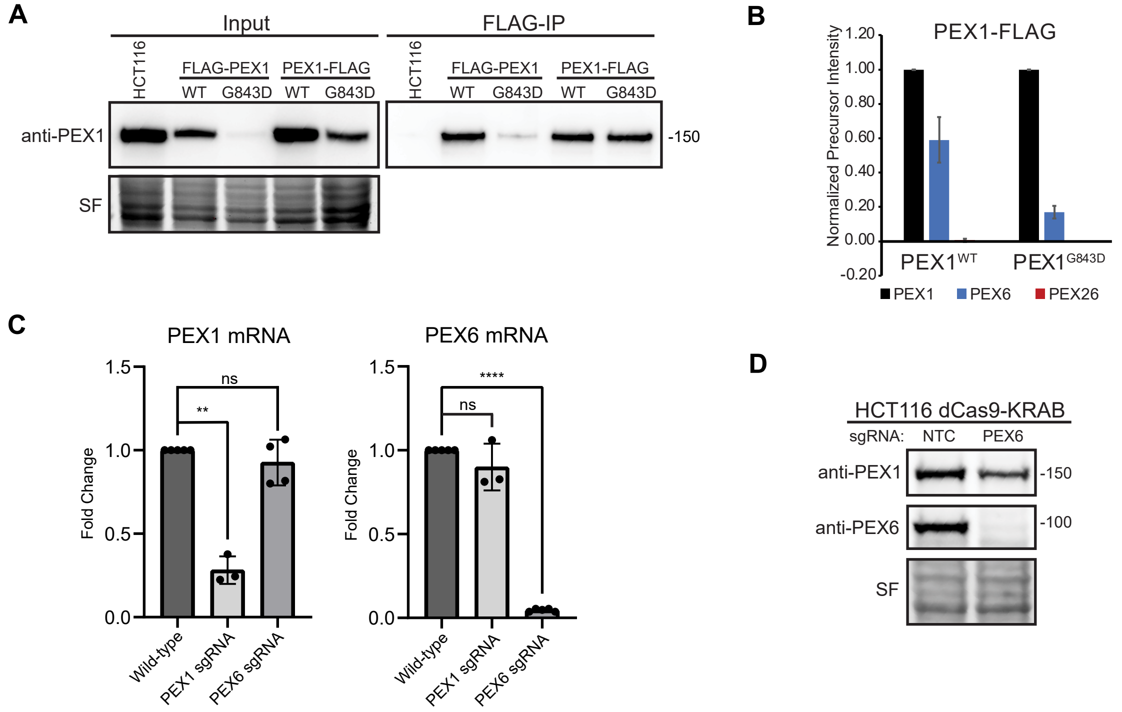

***Figure S5: Generation of CRISPRi PEX1$^{G843D}$ cells and candidate knockdown screen.***
A) Representative ClaI digest screen of CRISPR-Cas9 edited HCT116 clones for the G843D substitution. Clones A2 and B2 present homozygous clones with PEX1$^{G843D}$. B) Representative PEX1 immunoblot of CRISPRi PEX1$^{G843D}$ clones A2 and B2 shows reduced levels of PEX1 in both cell lines (PEX1 = 142 kDa, SF = StainFree total protein). C) RT-qPCR of CRISPRi cell lines with knockdowns of the candidate genes UBR5 and UBE2O identified by mass spectrometry.

### Figure S5

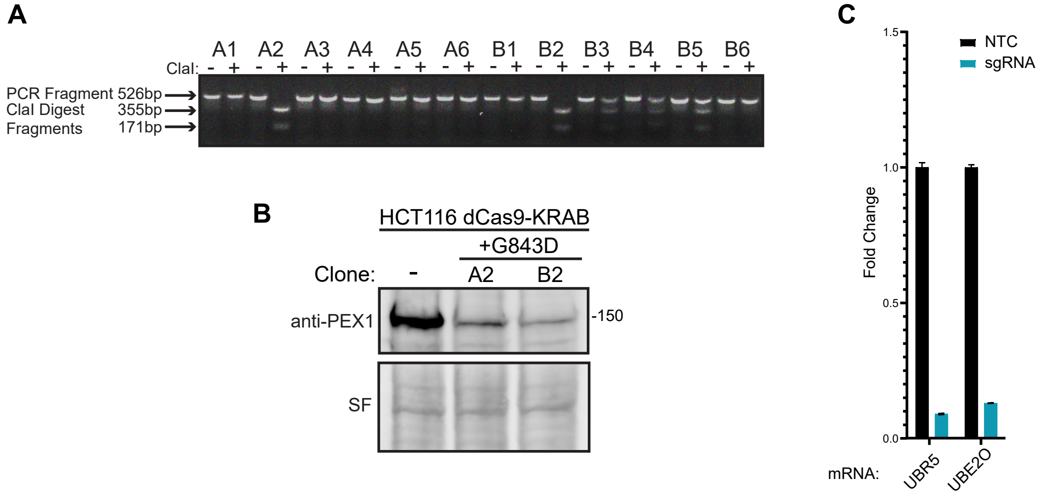
